# How many variables does WorldClim have, truly? Generative A.I. unravels the intrinsic dimension of bioclimatic variables

**DOI:** 10.1101/2023.06.12.544623

**Authors:** Russell Dinnage

## Abstract

The 19 standard bioclimatic variables available from the WorldClim dataset are some of the most used data in ecology and organismal biology. It is well known that many of the variables are correlated with each other, suggesting there are fewer than 19 independent dimensions of information in them. But how much information is there? Here I explore the 19 WorldClim bioclimatic variables from the perspective of the manifold hypothesis: that many high dimensional datasets are actually confined to a lower dimensional manifold embedded in an ambient space. Using a state-of-the-art generative probabilistic model (variational autoencoder) to model the data on a non-linear manifold reveals that only 5 uncorrelated dimensions are adequate to capture the full range of variation in the bioclimatic variables, with a clear data-driven separation between informative and redundant dimensions that eliminates arbitrary thresholds. I show that these 5 variables have meaningful structure and are sufficient to produce species distribution models (SDMs) nearly as good and in some ways better than SDMs using the original 19 bioclimatic variables. I have made the 5 synthetic variables available as a raster dataset at 2.5 minute resolution in an R package that also includes functions to convert back and forth between the 5 variables and the original 19.

## Introduction

The WorldClim dataset, often referred to as bioclim, is a widely utilized resource providing detailed global climate data suitable for a variety of ecological and biological applications (Fick and Hijmans 2017). The dataset comprises 19 bioclimatic variables, each of which represents a different aspect of annual trends, seasonality, and extreme or limiting environmental factors. These variables are second-order summaries derived from just three first-order monthly variables: minimum temperature, maximum temperature, and precipitation – via formulae computing means, extremes, ranges, and seasonal aggregates, providing more biologically meaningful summaries of environmental conditions than the raw monthly values alone. The set of 19 variables were selected over the period 1984-1996 as part of the development of BIOCLIM, the first species distribution modeling package (Booth et al. 2014), and this particular selection has remained the standard set of bioclimatic variables used in ecological studies ever since, despite lacking a formal analysis of their dimensionality.

The use of bioclim variables in species distribution modeling is well documented in the literature. These models frequently employ bioclim data to establish the ecological niche of a species, linking occurrences to the environmental conditions under which they are found (Elith and Leathwick 2009). One reason the dataset is so widely used, besides its easy and wide accessibility, is that equivalent variables have been estimated for past and future climates based on paleoclimatic reconstructions and future climate model projections respectively. This makes it straightforward to use species distribution models for prediction into the past and future, especially to predict shifts in species distributions under climate change scenarios, providing important information for conservation planning (Pearson et al. 2006). However, the reliability of SDM predictions can be sensitive to the choice and number of input variables, and high-dimensional correlated predictors can lead to spurious model performance (Fourcade et al. 2018).

However, a critical consideration when using bioclim variables is their inherent multicollinearity, due to the shared underlying data used in their computation. Techniques such as Principal Component Analysis (PCA), Variance Inflation Factor (VIF) analysis, and Regularization methods have been employed to address this issue, with the aim of producing more reliable and interpretable model outcomes (Dormann et al. 2013, Kriticos et al. 2014).

More generally, though each of the 19 bioclimatic variables measure an independently meaningful and potentially useful aspect of climate, when put together they likely contain considerable redundancy of information. Another way of putting this is that the ‘true’ or intrinsic dimension of the data is less than the total number of dimensions in the data, which can lead to problems for inference and prediction. For example, too many partially redundant variables can lead to variance inflation, difficulty in assessing the importance of individual variables, overfitting on noise, and decreased generalization ability in models (James et al. 2013). One way to deal with too many variables using dimension reduction methods such as Principle Components Analysis (PCA: (Pearson 1901)) to reduce the dimensionality of the dataset, a frequently used strategy when using bioclim data. Using PCA on bioclim can reduce the first 95% of variance into 10 orthogonal variables, but a closer look at the PCA results suggests that a linear dimension reduction method is inadequate. This is apparent from the fact that the cumulative variance explained by each additional dimension forms a smooth relationship (Figure 1), whereas we expect a discontinuous ‘jump’ or ‘elbow’ in this plot in the case of a true linear manifold (Cattell 1966). As I show below, a variational autoencoder provides a principled, data-driven alternative: latent dimensions either contribute to reconstruction (low posterior variance) or collapse to the prior (high variance), yielding an unambiguous separation without arbitrary thresholds.

**Figure 1.**
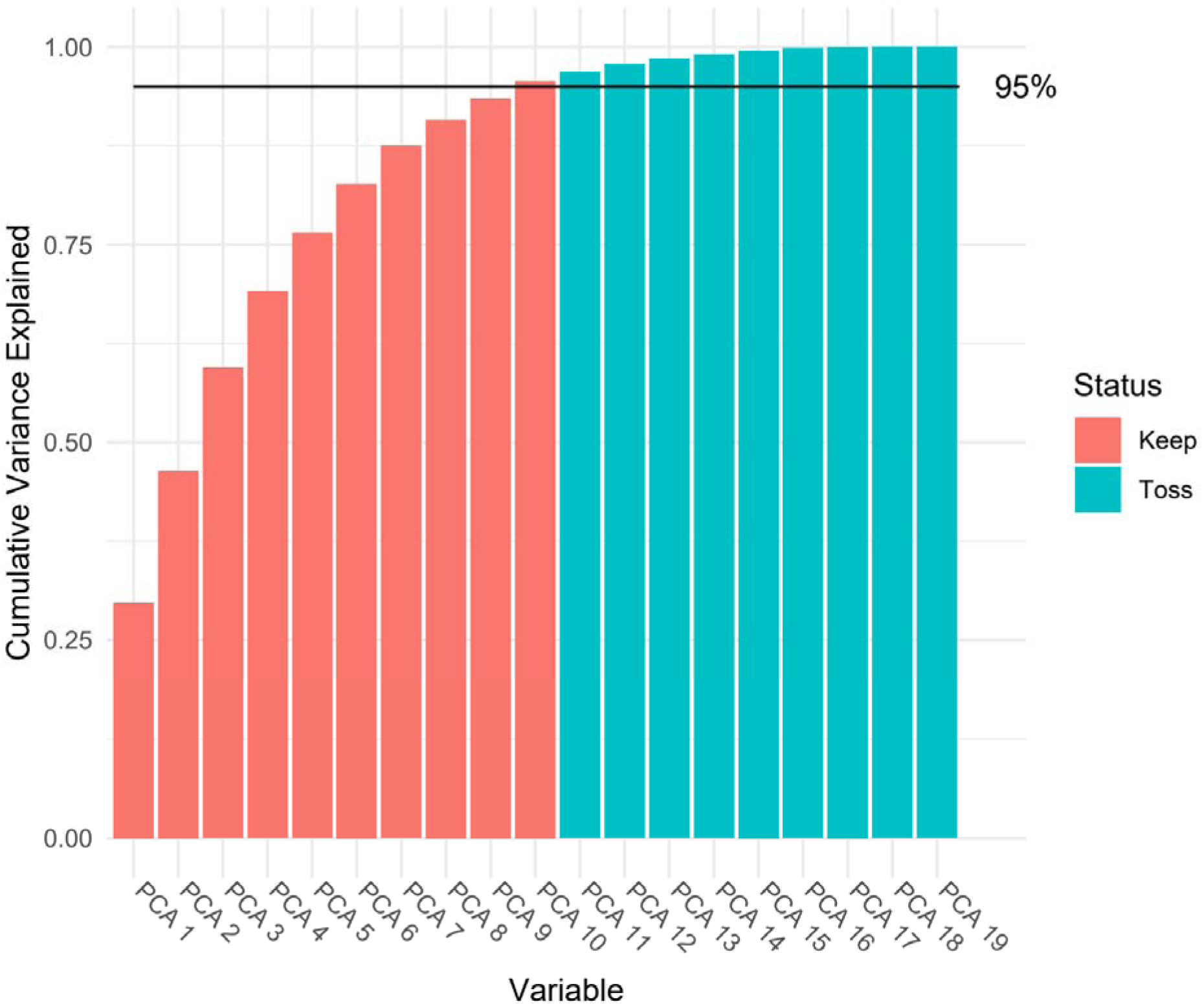
Cumulative variance explained by 19 PCA dimensions of bioclim variables. There is no obvious jump in the amount of new variance explained from one PCA dimension to the next, which would indicate a possible linear manifold. Otherwise, convention suggests arbitrarily keeping the PCA dimensions that cumulatively sum up to >95% of variance explained.

Manifold learning is an emerging concept rooted in the manifold hypothesis, which postulates that high-dimensional real-world data lie on or near a lower-dimensional manifold embedded within the high-dimensional space (Bengio et al. 2012). This hypothesis forms the basis for a wide variety of advanced dimension reduction and data analysis techniques, which include the new and exciting class of models known as deep generative learning. Classic PCA can also be considered a simple method of manifold learning, but where the manifold is assumed to be linear (e.g. a hyperplane). Here I explore the use of one deep generative model known as a variational autoencoder (VAE: (Kingma and Welling 2013)) for analysing bioclim variables, a simple but nevertheless useful example which I hope also demonstrates the exciting greater potential of VAEs and other generative models for ecology.

## Methods

### Variational Autoencoders

Variational autoencoders (VAEs) are a class of generative models that employ deep learning techniques for manifold learning (Kingma and Welling 2013). VAEs offer a probabilistic approach for representing complex, high-dimensional data in a lower-dimensional latent space.

These models are built on the principles of autoencoders, which are neural networks designed to learn a low dimensional representation of input data, through the use of a paired encoder and decoder model, that respectively encode the input into a latent space, and decode the latent space into the original input space (Hinton and Salakhutdinov 2006). VAEs introduce a layer of probabilistic inference, framing the encoding and decoding process as a probabilistic graphical model and utilizing Bayesian variational inference for learning and sampling (Kingma and Welling 2013).

Notably, VAEs encourage a specific structure on the latent space by assuming a simple Bayesian prior distribution for the latent space: an an independent Gaussian distribution for each latent variable. This promotes a continuous and well-structured latent space that facilitates the learning of high-dimensional manifold structures (Doersch 2016). Dai and Wipf 2019 and Zheng et al. 2023 recently showed that manifold dimensions can be identified by latent variables whose posterior distributions have a low variance.

VAEs have begun to receive attention in molecular biology for a variety of uses such as visualisation (Battey et al. 2020), estimating the probability density of sequence variants (Ziegler et al. 2023), generating candidate functional proteins (Hawkins-Hooker et al. 2020), and estimating reduced representations of complex metabolomics data useful for downstream predictive modeling. (Gomari et al. 2022), but they have yet to widely used in ecology. I hope this study, in addition to providing interesting new insights to a classical dataset used in ecology, can act as a concise introduction to VAEs for ecologists, and to promote their wider use, as their potential for ecology is substantial.

VAEs work by estimating two key functions, which are approximated using neural networks: an encoder, and a decoder.

The encoder, also known as the recognition or inference model, takes the input data and provides a probabilistic encoding in the latent space. Let’s say we have a dataset *X* = *x*_1_, *x*_2_, …, *x_n_* and we wish to encode it into a latent variable z. The encoder would provide us with a distribution *q_ϕ_* (*z*|*x*) over the latent variable, given the input data.

The decoder, also known as the generative or the likelihood model, takes a point in the latent space and generates a sample in the data space. It provides us with a distribution *p_θ_*(*x*|*z*) over the data given a latent variable.

The objective of VAE is to maximize the evidence lower bound (ELBO) on the log-likelihood of the data:

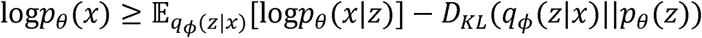

The first term on the right-hand side represents the expected reconstruction error – it encourages the VAE to reconstruct the input data accurately – and where the expectation is estimated using Monte Carlo sampling. The second term represents the Kullback-Leibler (KL) divergence between the approximate posterior *q_ϕ_* (*z*|*x*) and the prior *p_θ_*(*z*), which is typically chosen to be a independent normal distribution. This term acts as a regularization, encouraging the distribution of the latent variables to stay close to a unit Gaussian.

Training a VAE involves finding the parameters *θ* and *ϕ* that maximize the ELBO using stochastic gradient ascent, enabled by the reparameterization gradient, often known as the ”reparameterization trick” (Kingma and Welling 2013). The reparameterization trick allows a gradient calculation through the expectation in the ELBO, which is normally blocked by the non-differentiability of a random sampling operation. It works by reparameterizing the independent normal distribution *z* ∼ Normal(μ, ^2^) to *z* = μ + σ ⋅ ε, where ε ∼ Normal(0, 1). ε is sampled from a standard normal and then is injected into the model. Since the sampled stochastic noise is now an external input to the computational graph, the gradient no longer needs to pass through the sampling operation, allowing full backpropagation. After training a VAE, new data can be generated by sampling from the prior *p_θ_*(*z*), then passing these samples through the decoder.

A particularly useful aspect of VAEs for manifold learning is that maximizing the ELBO encourages the model to converge on a minimal set of ‘active’ latent variables to explain the data. A VAE learns an approximate Bayesian posterior distribution for each of the latent variables. Variables that contribute strongly to the reconstruction of the data tend to have low variance. On the other hand, any variables that are not important for reconstructing data will tend to have their variances converge to 1, consistent with the prior distribution, and facilitated by the flexibility of the decoder function, which can learn to ignore these high variance dimensions. Therefore, the number of active dimensions can be deduced by the number of variables with low variance posteriors (close to zero). Recent work has shown this to be a good estimate of the true manifold dimension of simulated data (Dai and Wipf 2019; Zheng et al. 2023).

The encoder and decoder neural networks were generated using the in-development dagnn R package (https://github.com/rdinnager/dagnn), which is designed to make specifying complex neural network architecture simple by specifying a directed acyclic graph (DAG) using R formula syntax.

**Table 1.** Glossary of Variational Autoencoder related terms.

| Term | Description |
| --- | --- |
| Generative Models | A class of models that learn the full process generating a dataset, not just how to predict an outcome. Unlike models that only learn input-output relationships (e.g. a GLM predicting species presence from climate), generative models can also create (simulate) new realistic data points, because they capture the overall structure of the data. |
| Autoencoder (AE) | A neural network that learns to compress data into a small number of variables (encoding) and then reconstruct the original data from them (decoding). The compressed representation captures the most important patterns. A deterministic precursor to VAEs that lacks the probabilistic framework. |
| Variational Inference | A method for approximating complex probability distributions that would be too expensive to compute exactly. In VAEs, it is used to estimate the most likely values of the latent (hidden) variables given the observed data, analogous to how Bayesian methods estimate posterior distributions but using optimisation rather than sampling (e.g. MCMC). |
| Kullback-Leibler Divergence (KL) | A measure of how much one probability distribution differs from another. In VAEs, it measures how far the learned latent variable distributions deviate from the assumed prior (a standard normal distribution), acting as a penalty that keeps the latent space well-structured and prevents the model from encoding too much specific |
|  | information about individual data points. |
| Regularization | Any technique that constrains a model during training to prevent it from fitting noise in the data (overfitting). In VAEs, the KL divergence term acts as a regularizer: it penalises latent distributions that deviate far from the prior, encouraging the model to use only as many latent dimensions as are truly needed to represent the data. |
| Evidence Lower Bound (ELBO) | The objective function maximised during VAE training. It balances two competing goals: (1) accurately reconstructing the input data (reconstruction loss) and (2) keeping the latent variable distributions simple and close to the prior (KL divergence). The name comes from the fact that maximising it approximates maximising the probability of the observed data (the “evidence”) under the model. |
| Reparameterization Trick | A mathematical technique that enables gradient-based training of VAEs despite the random sampling step. Instead of sampling latent values $z$ directly from a learned distribution, we compute $z = \text{mean} + \text{standard\_deviation} \times \text{noise}$, where the noise is drawn from a fixed standard normal distribution. This separates the randomness from the learnable parameters (mean and standard deviation), allowing the training algorithm to compute how to adjust them. |
| Manifold | A smooth, lower-dimensional surface embedded within a higher- |
|  | dimensional space. For example, the surface of the Earth is a 2-dimensional manifold in 3-dimensional space. The manifold hypothesis proposes that complex high-dimensional data (such as 19 bioclim variables) actually lie on or near such a lower-dimensional surface, meaning fewer variables are needed to describe the data than the number of measured variables. |
| Latent Variables / Latent Space | Hidden (unobserved) variables that a model infers from the data. In a VAE, the latent space is the low-dimensional representation that the encoder produces and the decoder reconstructs from. Each point in the latent space corresponds to a particular combination of the original variables. “Active” latent variables are those the model actually uses; inactive ones are ignored by the decoder. |
| Posterior Distribution | In Bayesian statistics, the probability distribution of unknown quantities (e.g. latent variables) after accounting for the observed data. In a VAE, for each data point the encoder estimates a posterior distribution over the latent variables, characterised by a mean and variance for each latent dimension. |

### Model Training

I trained a VAE model coded in R with the torch package (Falbel and Luraschi 2022) on 19 global bioclimatic variables at 2.5 minute resolution, for a total of 12,532,612 data points, one for each global grid-cell. Full code to reproduce the model and all analyses in this study is available at: https://github.com/rdinnager/bioclim_intrinsic_dimension. I allowed for a maximum of 64 latent dimensions. The bioclim variables were standardised to zero mean and unit standard deviation before model input. Key hyperparameters included: 16 initial latent dimensions, a 3-layer encoder and decoder with skip connections, and the trainable precision parameter ‘gamma’ from Dai and Wipf (2019). The model was trained for 1000 epochs using the Adam optimiser with a one-cycle learning rate schedule (maximum learning rate 0.005). The VAE was trained using the procedure recommended in (Dai and Wipf 2019), which includes as trainable ‘gamma’ parameter. ‘Gamma’ in this case is the variance of the Gaussian distribution used to calculated the log likelihood of the VAE reconstruction. This gamma parameter tends to decrease during training as the reconstruction gets better, which has the effect of increasing the weighting of the reconstruction loss relative to the KL loss as the training goes on. Dai and Wipf (2019) showed this leads to a much better reconstruction whilst still maintaining a well-structured latent space.

### Interpretation

To explore how to interpret the manifold variables extracted by the VAE, I looked at how the they were related to the original 19 variables, as well as how they were represented within the world’s major biomes, using data from Dinerstein et al. (2017).

### Comparing Species Distribution Models

I ran three sets of species distribution models (SDMs) using the *ENMTools* R package (Warren et al. 2021): 1) SDMs using all 19 original bioclim variables; 2) SDMs using the first 10 PCA axes of the 19 bioclim variables (computed as part of this study, explaining ∼95% of total variance; Figure 1); and 3) SDMs using the 5 manifold variables estimated in this study. This was done using three different SDM methods: Random Forest, GLM, and the Bioclim model. The three sets of models were run on occurrence data taken from (Elith et al. 2020), which is available in the R package *disdat.* I used 5 out of the 6 datasets provided by Elith et al. (2020) (the 6th proved too large to use easily in the parallelized pipeline developed for this study). The 5 datasets contained a total of 206 species from 5 different areas:

1) Australian Wet Tropics (Queensland, Australia): 20 bird species, and 20 plant species
2) Ontario, Canada: 30 bird species
3) New South Wales, Australia: 7 bat species, 10 bird species, 29 plant species, 8 small reptile species
4) New Zealand: 52 vascular plant species
5) Switzerland: 30 tree species

For each species and model combination I ran 10 replicate SDMs using ENMTools, holding out a random 10% of the occurrence points in each replicate. I then calculated the Reciever-Operator Curve Area Under the Curve metric (AUC) of model fit on model predictions for the held out 10%. I also calculated the True Skill Statistic (TSS) as an alternative measure of model fit.

Additionally, I used the *ENMTools*’ env.evaluate() function to calculate AUC and TSS in environmental space. This uses latin hypercube sampling in environmental space to sample background points which can be used in the same way that geographic background points are used to estimate the ability of the model to predict presence or absence in environmental space.

To compare the bioclim manifold variables and the bioclim PCA variables I calculated a metric *Δ*AUC for each replicate of each species and model, which is the different between the AUC metric of the focal SDM (manifold or PCA) and the baseline model, which in this case is the corresponding model run on all 19 original bioclim variables.

I report 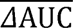 in the main text, which is the average *Δ*AUC over all replicates, models and species. In order to test whether this value was significantly different than zero, I fit a Linear Mixed Model (LMM) using the *lme4* R package, including the variable type (manifold vs. PCA) as a fixed effect and species, area (e.g. New Zealand vs. Canada), and model as random effects. The reported 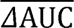 values are the estimated marginal means from this model (calculated using the *emmeans* R package). The same was done for the True Skill Statistic.

Lastly, the whole above procedure was repeated but using a simple spatial block technique (in which the study area is partitioned into geographic blocks and entire blocks are withheld for testing, rather than random individual points) for generating the hold-out sample instead of a simple random sample. For this I used the spatial block sampling option in ENMTools model fitting function. This tests whether models can generalise to new geographic areas, a critical requirement for SDMs used in conservation planning.

## Results

### The Instrinsic Dimension of WorldClim

After model training, examining the posterior distribution of the 64 latent dimensions revealed only 5 had low variance (Figure 2), suggesting an intrinsic dimensionality of 5. Using only these 5 variables to reconstruct the original data through the trained decoder, the reconstruction error is extremely low, with a mean squared error of reconstruction of 0.0036 (on the standardised variables, so in units of standard deviations).

**Figure 2.**
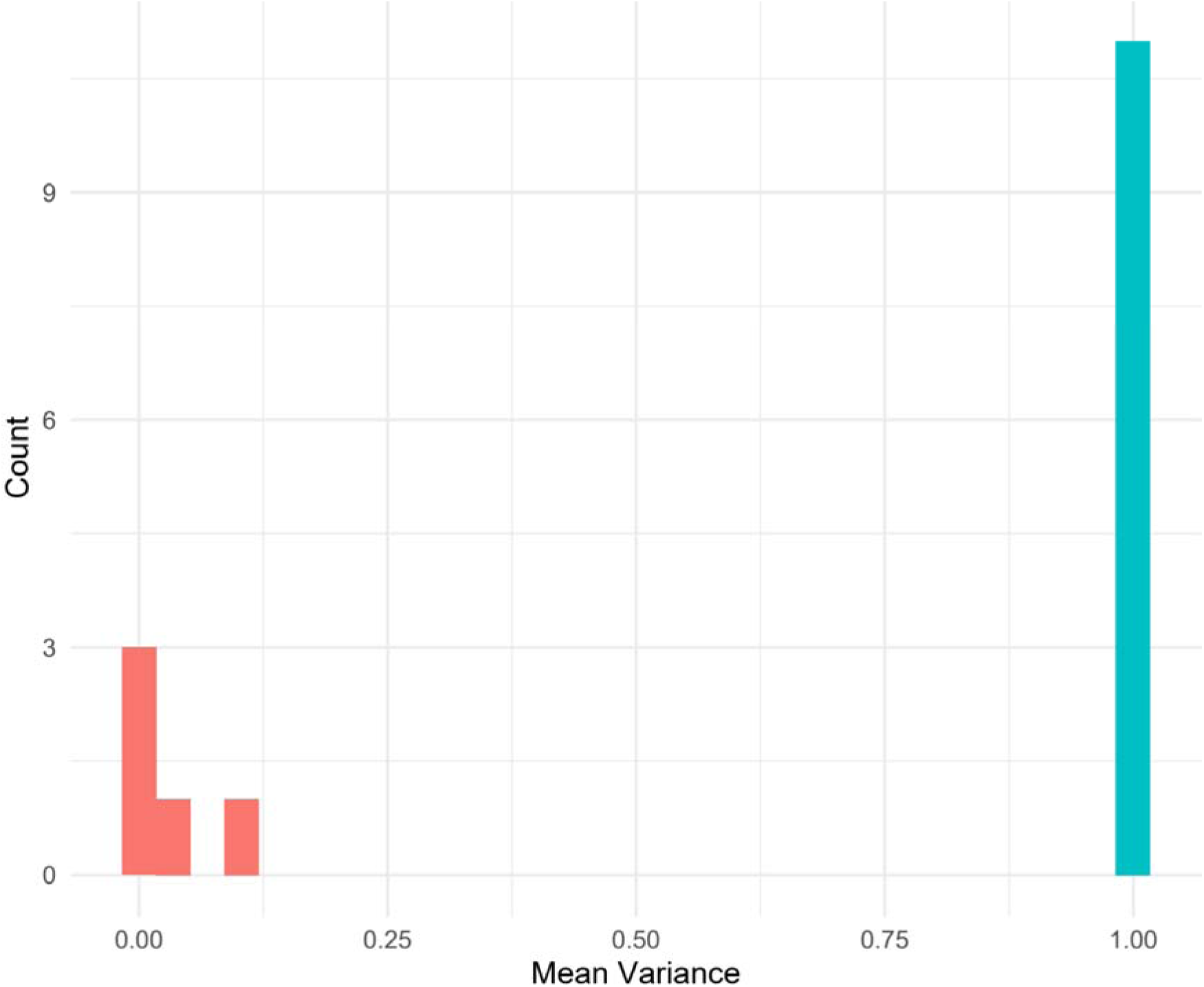
The mean variances along each variational autoencoder latent dimension (there were 16 altogether). “Mean Variance” refers to the approximate posterior variance for each latent dimension, averaged across all ∼12.5 million data points. All dimensions except for 5 had variance near or equal to 1, matching the prior distribution. The remaining 5 had variances close to zero. This is indicative that these are ‘manifold’ dimensions: they are necessary to reconstruct the full data. All other dimensions are just noise that the decoder network learns to ignore.

Using SDM benchmark data from (Elith et al. 2020), and three different SDM methods (random forest, glm, and bioclim) from the ENMTools R package (Warren et al. 2021), I additionally tested whether these 5 ‘latent’ variables are useful by comparing species distribution models fit with them with models fit on all 19 original variables, as well as the first 10 PCA dimensions as a comparison. For the 5 manifold dimensions and the PCA axes I calculated the model performance using 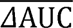, defined as the average difference in AUC between the focal model and the corresponding model using all 19 original variables, averaged over all species and methods tested. I find the 5 variables perform slightly less well (but not significantly) at predicting held-out test samples in geographic space (i.e. at spatially withheld occurrence locations) according to AUC (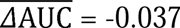 [CI: -0.076, 0.002]; -0.02 for PCA [CI: -0.058, 0.019]; Figure S1) but perform considerably better at predicting them in environmental space (i.e. at background points sampled across the range of environmental conditions rather than geographic locations; Warren et al. 2018) ( 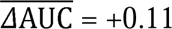 [CI: 0.071, 0.15] , +0.047 for PCA [CI: 0.0096, 0.087]; Figure S1) (Warren et al. 2018). The same analysis using the true skill statistic (TSS) instead of AUC gave similar qualitative results (Figure S2). I suspected the slightly worse performance in geographic space was because with fewer variables, models could not overfit as strongly to approximate spatial structure in the occurrence data. To test this I repeated the SDMs using a simple spatial block method for holding out test samples. This confirmed my suspicion, with the reduction in AUC for spatially held-out test data being much smaller and again, when evaluated in geographic space (i.e. predicting at spatially withheld locations), the confidence interval for the mean delta-AUC included zero, indicating no statistically significant difference from the full 19-variable models (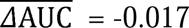 [CI: -0.043, 0.0081]; Figure S3), but still much improved in environmental space ( 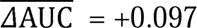 [CI: 0.071, 0.12]; Figure S3).

The 5 manifold dimensions learned by the VAE in this study have been provided in an R package (https://github.com/rdinnager/biocman), which also provides function for converting between the 5 manifold variables and the original 19 bioclimatic variable (e.g. functions for encoding and decoding).

## Discussion

### Interpreting the Manifold Dimensions

Examining the 5 active variables shows that each has a unique global pattern (Figure 3) and link to the original 19 variables (Figures S4-S8). Three of the variables admit a fairly straightforward interpretation, BIOCMAN1, BIOCMAN2, and BIOCMAN4 – their values have a mostly monotonic relationships with the original 19, and their global distribution maps show clear links to important global climate factors, BIOCMAN1 being elevation, BIOCMAN2 being areas supporting rainforest, including temperate rainforest, and BIOCMAN4 being latitude. The remaining two are less straightforward, with some relationships with both the original 19 variables being unimodal or ‘U-shaped’. On the other hand, BIOCMAN5 appears to clearly differentiate areas with monsoonal climates with high values. BIOCMAN3 has the least clear interpretation but globally low values seem to be associated with arid and semi-arid ecoregions (Figure 1; Figure S6).

**Figure 3.**
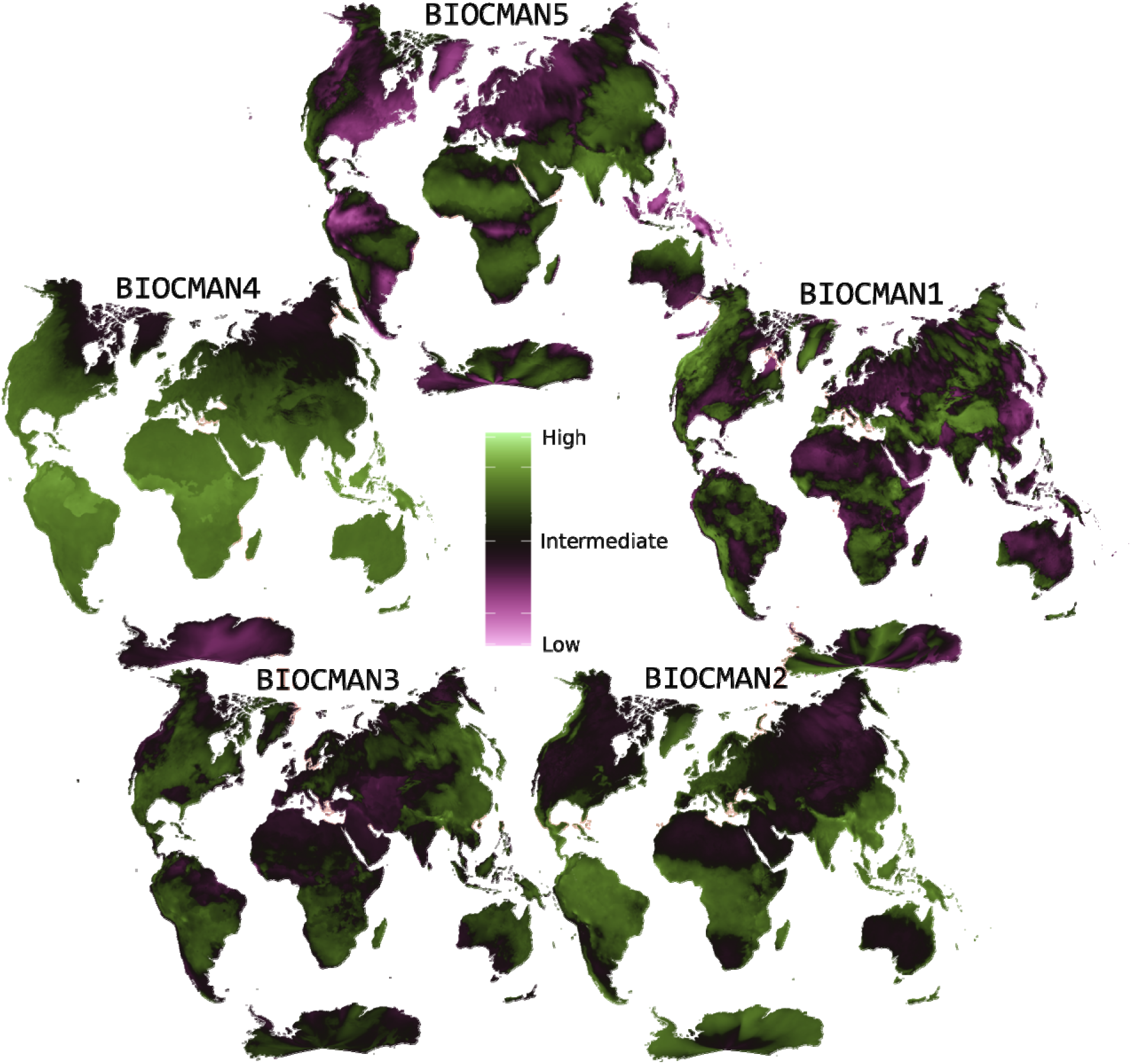
Global maps of the 5 VAE manifold variables. Most variables appear to be imperfect but reasonable approximations to known global climate factors. BIOCMAN1 looks like elevation. BIOCMAN2 is where rainforest exists (including temperate rainforest). BIOCMAN3 has the least clear pattern, but low values seem to be associated with arid and semi-arid places. BIOCMAN4 is close to a map of latitude, and BIOCMAN5 is high where there are monsoonal climates. Maps are presented in Danseiji V projection, which has low distortion in terrestrial areas at the cost of high distortion in the oceans (Kunimune 2020). High resolution versions with full scale bars can be found in Appendix S1: Figures S9–S13.

For the application of species distribution modeling and other ecological models where interpretation of a model’s coefficients might be of interest, the issue of challenging interpretability can be mitigated using the decoder function, which allows a straightforward way to convert the manifold variables back to the original data space. In this way they are no less interpretable than using a transformation to make a variable better conform to model assumptions (e.g. log or logistic transformations to reduce covariate skew). A simple procedure to interpret model coefficients when using bioclim manifold variables as predictors is to output predictions of the model across a range of manifold variable values of interest, then plot these predictions against the original 19 variables after back-transforming the manifold variables using the decoder. This then places the model prediction back into the context of full bioclimatic variation. For instance, if an SDM shows a strong effect of BIOCMAN1, the decoder reveals this corresponds primarily to variation in temperature-related variables (e.g. BIO1, Annual Mean Temperature), allowing interpretation in terms of familiar bioclimatic variables. This approach is analogous to examining PCA loadings, which map principal components back to original variables. However, PCA loadings are single numbers, a direct consequence of the linear assumption, making them easy to tabulate but potentially misleading when the true relationships are non-linear. The VAE decoder relaxes this constraint, capturing non-linear mappings that may require visualization rather than a single coefficient (see Supplementary Material: Figures S4–S8), but in return provides a more faithful representation of the underlying data structure.

Though many ecologists probably suspected that bioclim had fewer intrinsic dimensions than 19, including myself, it is perhaps surprising that the full panoply of bioclimatic variation on Earth can be reconstructed with as few as 5 variables. It is also remarkable that these 5 variables have close correspondence with known sources of climate variation such as latitude, elevation, and the monsoon. This speaks to the potential power of generative A.I. methods in ecology. In this case the physical and biological processes of the earth encode themselves in global climate variation, which this VAE was then able to decode, thanks to its ability to learn flexible non-linear functions of a simplified underlying space.

The existence of this low-dimensional manifold has a concrete explanation. The 19 bioclim variables are derived from just three independent monthly variables – minimum temperature, maximum temperature, and precipitation across 12 months (36 values in total) – through formulae that compute means, extremes, ranges, and seasonal aggregates (Booth et al. 2014). These derivation formulae impose deterministic algebraic relationships among the 19 variables, confining them to a non-linear surface within the 19-dimensional ambient space. The 36 underlying monthly values are themselves highly correlated through seasonal cycles, latitude effects, and temperature–precipitation coupling, so their effective dimensionality is considerably less than 36. The bioclim formulae further compress this variation, consistent with the 5-dimensional manifold the VAE recovers. Importantly, the manifold dimensions do not simply invert the bioclim derivation formulae to recover monthly values; rather, they capture non-linear combinations reflecting physical climate processes like elevation gradients and monsoon dynamics. One might ask why not use the monthly variables directly – the answer is that the bioclim summaries encode biologically meaningful aggregates (e.g. temperature of the warmest quarter, precipitation seasonality) that are more directly relevant to species’ physiological tolerances, and the VAE’s bidirectional encoder–decoder preserves this biological relevance while removing redundancy.

A key methodological contribution of this work is therefore a principled, automatic approach to dimensionality selection. Unlike PCA, where the number of components to retain requires subjective judgment (e.g. variance thresholds, scree plot “elbows”), the VAE’s regularization structure produces a clear binary separation between active and inactive dimensions (Figure 2), eliminating arbitrary choices entirely. This yielded 5 uncorrelated variables that perform just as well as the original 19 in at least one downstream application (SDMs), and in some ways better. If even just for this automatic dimensionality selection, VAE methods are currently under-used in ecology.

A potential limitation of this study is that standard VAEs are non-identifiable: different training runs may learn different rotations of the same latent subspace (Locatello et al. 2019). To address this concern, I conducted an identifiability analysis by training 16 identifiable VAE (iVAE; Khemakhem et al. 2020) models using latitude and longitude, encoded as sinusoidal angle embeddings (see Supplementary Material), as auxiliary conditioning information, of which 9 converged to the same 5 active latent dimensions as the original VAE. Using the Mean Correlation Coefficient (MCC) with Procrustes alignment to compare learned representations across runs, I found that the manifold subspace is highly reproducible (mean Procrustes MCC = 0.84 across all pairwise comparisons). The original (unconditional) VAE showed strong agreement with the iVAE runs (mean Procrustes MCC = 0.81) and clustered among them in a hierarchical clustering analysis rather than as an outgroup, confirming that the 5-dimensional manifold structure is a genuine feature of the bioclimatic data rather than an artifact of a particular training run (see Supplementary Material: Identifiability of VAE Manifold Dimensions for full details; Figure S14). Not all dimensions were equally reproducible: the best-matched dimension pair across runs had a mean absolute correlation of 0.96, while the weakest had 0.65, suggesting that some manifold dimensions capture more robust climatic gradients than others (Figure S14D).

The VAE-based variable selection approach demonstrated here has potential applications beyond bioclimatic data. The ability to automatically determine intrinsic dimensionality from data, without requiring ad hoc thresholds, could prove valuable wherever ecologists work with large suites of correlated environmental variables—for example, remote sensing spectral bands, soil chemistry profiles, or oceanographic variables. Recent work has demonstrated that VAEs can compress large climate reanalysis datasets by factors of several hundred while retaining essential structure (Han et al. 2025), confirming that climate data generally occupies a low-dimensional manifold. More broadly, deep learning approaches are increasingly being applied to species distribution modeling (Kellenberger et al. 2026), and the manifold variables developed here can serve as inputs to such models. The NicheFlow framework (Dinnage 2024) extends the present work by using normalizing flows to build generative models of species’ environmental niches directly in manifold space, enabling both niche characterization and novel environmental prediction.

The present analysis was conducted at 2.5 arc-minute resolution. Preliminary experiments at 5 arc-minute resolution also recovered 5 active manifold dimensions, suggesting that the intrinsic dimensionality of bioclimatic variation is relatively robust to spatial resolution within this range. A formal multi-resolution sensitivity analysis using the iVAE framework with warm-start training across resolutions is an important avenue for future work. Previous studies have generally found that changing spatial resolution affects the magnitude but not the qualitative nature of species–environment relationships (Merkenschlager et al. 2023), consistent with our observation that the manifold structure is preserved across the resolutions examined.

## References

Battey, C. J., G. C. Coffing, and A. D. Kern. 2020. Visualizing population structure with variational autoencoders. G3: Genes|Genomes|Genetics 11.

Bengio, Y., A. C. Courville, and P. Vincent. 2012. Representation Learning: A Review and New Perspectives. IEEE Transactions on Pattern Analysis and Machine Intelligence 35:1798–1828.

Booth, T. H. 2022. Checking bioclimatic variables that combine temperature and precipitation data before their use in species distribution models. Austral Ecology.

Booth, T. H., H. Nix, J. R. Busby, and M. F. Hutchinson. 2014. bioclim: the first species distribution modelling package, its early applications and relevance to most current MaxEnt studies. Diversity and Distributions 20.

Cattell, R. B. 1966. The Scree Test For The Number Of Factors.. Multivariate behavioral research 1 2:245–76.

Dai, B., and D. P. Wipf. 2019. Diagnosing and Enhancing VAE Models. ArXiv abs/1903.05789.

Dinerstein, E., D. P. Olson, A. R. Joshi, C. Vynne, N. D. Burgess, E. D. Wikramanayake, N. R. Hahn, S. Palminteri, P. Hedao, R. F. Noss, M. Hansen, H. Locke, E. C. Ellis, B. S. Jones, C. V. Barber, R. Hayes, C. F. Kormos, V. G. Martin, E. Crist, W. Sechrest, L. Price, J. E. M. Baillie, D. E. Weeden, K. F. Suckling, C. Davis, N. C. Sizer, R. Moore, D. Thau, T. Birch, P. V. Potapov, S. Turubanova, A. Tyukavina, N. de Souza, L. Pintea, J. C. Brito, O. A.-ar-R. Llewellyn, A. G. Miller, A. Patzelt, S. A. Ghazanfar, J. R. Timberlake, H. Klöser, Y. Shennan Farpón, R. Kindt, J.-P. B. Lillesø, P. van Breugel, L. Graudal, M. Voge, K. F. Al-Shammari, and M. Saleem. 2017. An Ecoregion-Based Approach to Protecting Half the Terrestrial Realm. Bioscience 67:534–545.

Doersch, C. 2016. Tutorial on Variational Autoencoders. ArXiv abs/1606.05908.

Dinnage, R. 2024. NicheFlow: Towards a foundation model for Species Distribution Modelling. bioRxiv. 10.1101/2024.10.15.618541.

Dormann, C. F., J. Elith, S. Bacher, C. M. Buchmann, G. Carl, G. Carré, J. R. G. Márquez, B. Gruber, B. Lafourcade, P. J. Leitão, T. Münkemüller, C. J. McClean, P. E. Osborne, B. Reineking, B. Schröder, A. K. Skidmore, D. Zurell, and S. Lautenbach. 2013. Collinearity: a review of methods to deal with it and a simulation study evaluating their performance. Ecography 36:27–46.

Elith, J., C. H. Graham, R. Valavi, M. Abegg, C. Bruce, A. J. Ford, A. Guisan, R. J. Hijmans, F. Huettmann, L. G. Lohmann, B. A. Loiselle, C. C. Moritz, J. M. C. Overton, A. T. Peterson, S. J. Phillips, K. S. Richardson, S. E. Williams, S. K. Wiser, T. Wohlgemuth, and N. E. Zimmermann. 2020. Presence-only and Presence-absence Data for Comparing Species Distribution Modeling Methods. Biodiversity Informatics 15:69–80.

Elith, J., and J. R. Leathwick. 2009. Species Distribution Models: Ecological Explanation and Prediction Across Space and Time. Annual Review of Ecology, Evolution, and Systematics 40:677–697.

Falbel, D., and J. Luraschi. 2022. torch: Tensors and Neural Networks with ’GPU’ Acceleration.

Fick, S. E., and R. J. Hijmans. 2017. WorldClim 2: new 1 km spatial resolution climate surfaces for global land areas. International Journal of Climatology 37.

Fourcade, Y., A. G. Besnard, and J. Secondi. 2018. Paintings predict the distribution of species, or the challenge of selecting environmental predictors and the modelling procedure. A reply to Titley et al. Global Ecology and Biogeography 27:245–256.

Gomari, D. P., A. Schweickart, L. Cerchietti, E. Paietta, H. Fernandez, H. Al-Amin, K. Suhre, and J. Krumsiek. 2022. Variational autoencoders learn transferrable representations of metabolomics data. Communications Biology 5.

Han, T., Z. Chen, S. Guo, W. Xu, W. Ouyang, and L. Bai. 2025. Climate science data can be compressed efficiently by dual-stage extreme compression with a variational auto-encoder transformer. Communications Earth & Environment 6:955.

Hawkins-Hooker, A., F. Depardieu, S. Baur, G. Couairon, A. Chen, and D. Bikard. 2020. Generating functional protein variants with variational autoencoders. PLoS Computational Biology 17.

Hinton, G. E., and R. Salakhutdinov. 2006. Reducing the Dimensionality of Data with Neural Networks. Science 313:504–507.

James, G., D. Witten, T. Hastie, and R. Tibshirani. 2013. An Introduction to Statistical Learning. Springer, New York.

Kellenberger, B., K. Winner, and W. Jetz. 2026. The Performance and Potential of Deep Learning for Predicting Species Distributions. Global Ecology and Biogeography 35:e70184.

Khemakhem, I., D. Kingma, R. Monti, and A. Hyvärinen. 2020. Variational autoencoders and nonlinear ICA: A unifying framework. Proceedings of the 23rd International Conference on Artificial Intelligence and Statistics (AISTATS), PMLR 108:2207–2217.

Kingma, D. P., and M. Welling. 2013. Auto-Encoding Variational Bayes. CoRR abs/1312.6114.

Kriticos, D. J., Jaroık Vojtěch, and N. Ota. 2014. Extending the suite of bioclim variables: a proposed registry system and case study using principal components analysis. Methods in Ecology and Evolution 5.

Kunimune, J. H. 2020. Minimum-error world map projections defined by polydimensional meshes. International Journal of Cartography 7:78–99.

Locatello, F., S. Bauer, M. Lucic, G. Rätsch, S. Gelly, B. Schölkopf, and O. Bachem. 2019. Challenging common assumptions in the unsupervised learning of disentangled representations. Proceedings of the 36th International Conference on Machine Learning (ICML), PMLR 97:4114–4124.

Merkenschlager, C., F. Bangelesa, H. Paeth, and E. Hertig. 2023. Blessing and curse of bioclimatic variables: A comparison of different calculation schemes and datasets for species distribution modeling within the extended Mediterranean area. Ecology and Evolution 13:e10553.

Pearson, K. 1901. LIII. On lines and planes of closest fit to systems of points in space. Philosophical Magazine Series 1 2:559–572.

Pearson, R. G., W. Thuiller, M. B. Araújo, E. Martínez Meyer, L. Brotóns, C. J. McClean, L. Miles, P. Segurado, T. P. Dawson, and D. C. Lees. 2006. Model based uncertainty in species range prediction. Journal of Biogeography 33.

Warren, D. L., L. J. Beaumont, R. Dinnage, and J. B. Baumgartner. 2018. New methods for measuring ENM breadth and overlap in environmental space. Ecography.

Warren, D. L., N. J. Matzke, M. Cardillo, J. B. Baumgartner, L. J. Beaumont, M. Turelli, R. E. Glor, N. A. Huron, M. V. P. Simões, T. L. Iglesias, J. C. Piquet, and R. Dinnage. 2021. ENMTools 1.0: an R package for comparative ecological biogeography. Ecography.

Zheng, Y., T. He, Y. Qiu, and D. P. Wipf. 2023. Learning Manifold Dimensions with Conditional Variational Autoencoders. ArXiv abs/2302.11756.

Ziegler, C., J. Martin, C. Sinner, and F. Morcos. 2023. Latent generative landscapes as maps of functional diversity in protein sequence space. Nature Communications 14.

