## Supporting Information for "How many variables does WorldClim have, truly? Generative A.I. unravels the intrinsic dimension of bioclimatic variables"

### Supplementary Material

#### Figures

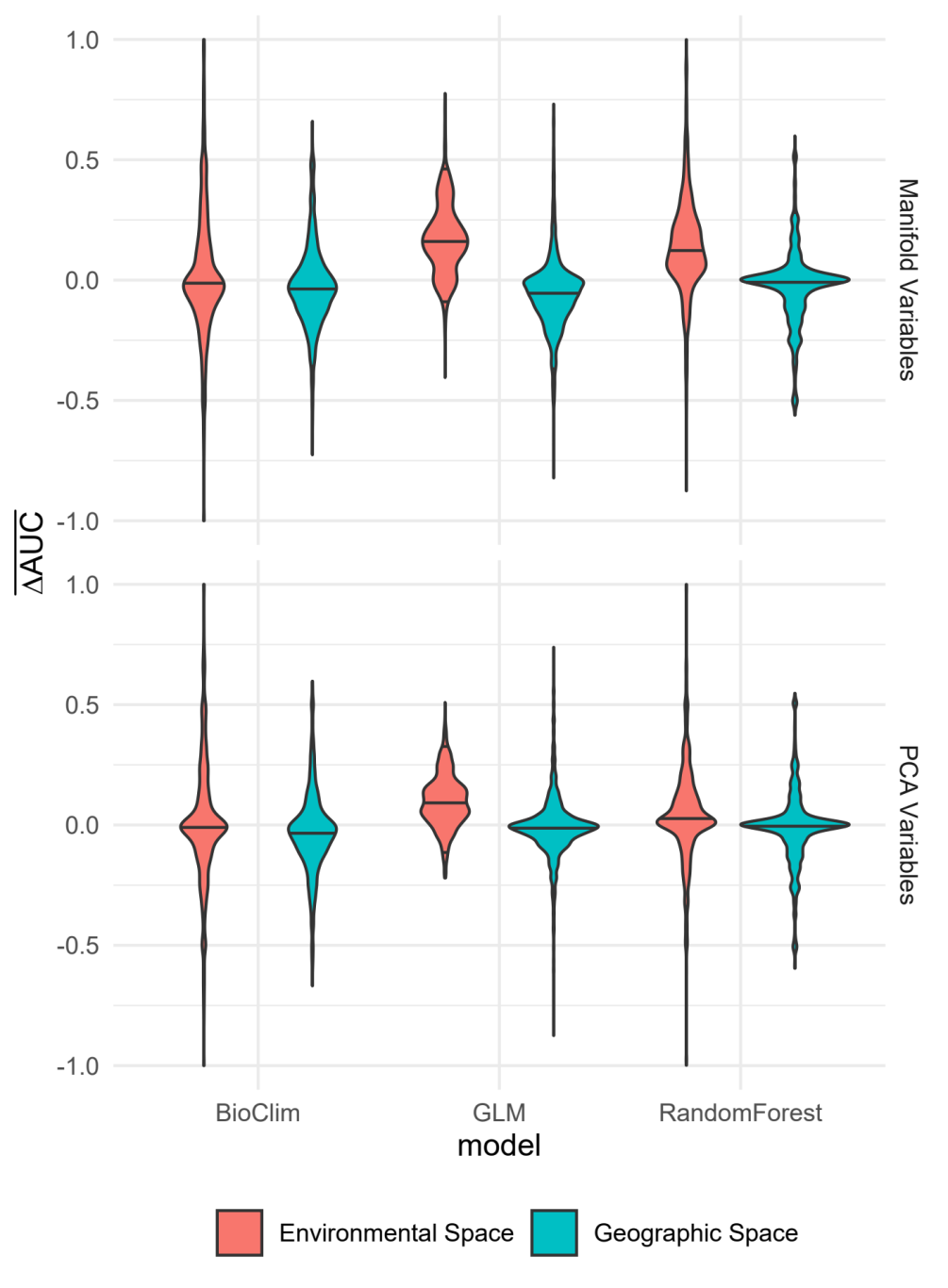

**Figure S1.** Violin plot of random hold-out $\overline{\Delta AUC}$, by model type (X axis), Environmental vs. Geographic space (colour) and whether the models used the manifold or the PCA variables (top and bottom panels). Width of the violins is proportional to the density of points at the value on the y axis. Horizontal black lines represent the 2.5%, 50%, and 97.5% quantiles.  $\overline{\Delta AUC}$ is the difference in AUC between the focal model and the corresponding model using the original 19 bioclim variables, calculated on a 10% random hold-out sample.

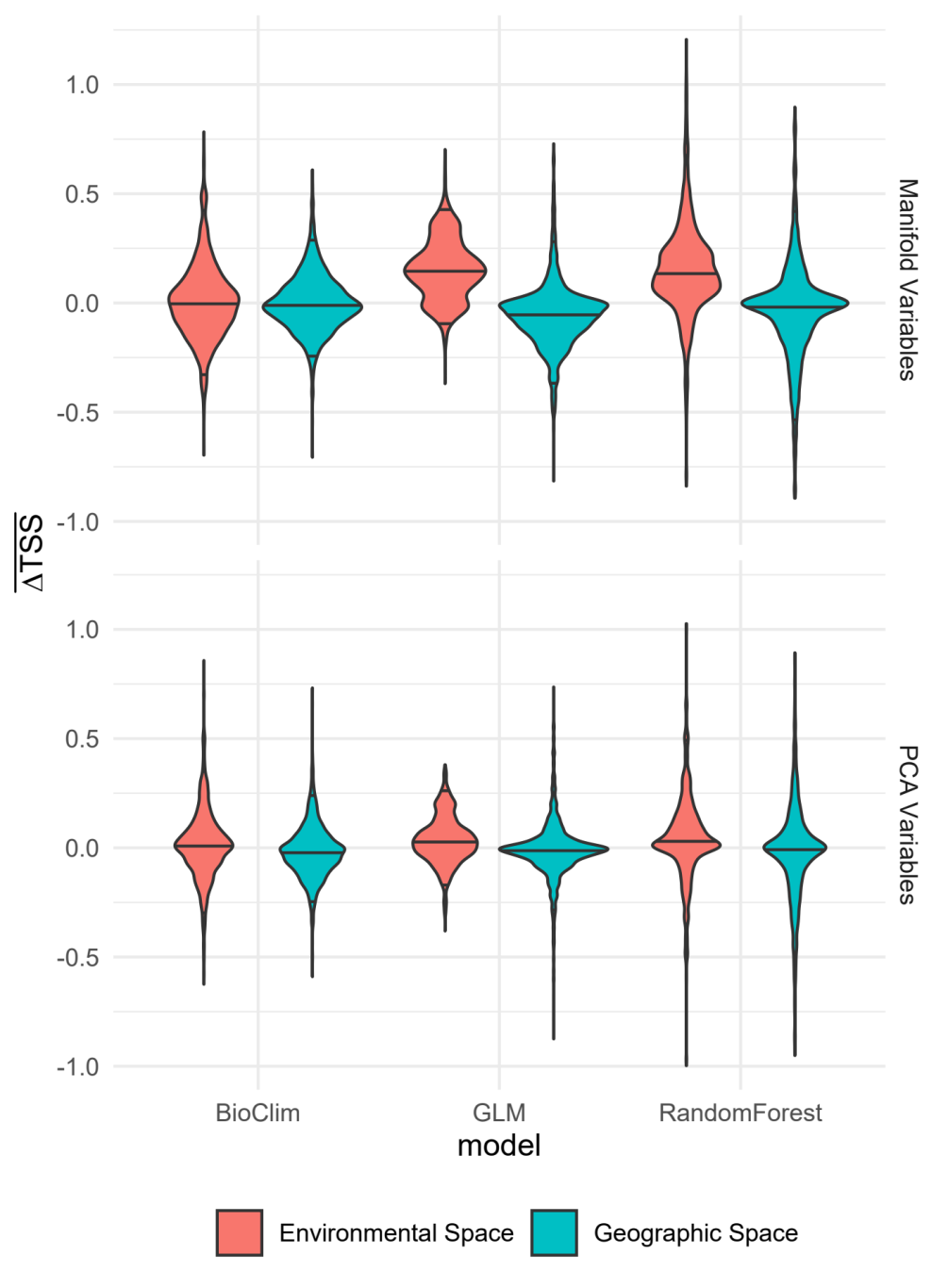

**Figure S2.** Violin plot of random hold-out $\overline{\Delta TSS}$, by model type (X axis), Environmental vs. Geographic space (colour) and whether the models used the manifold or the PCA variables (top and bottom panels). Width of the violins is proportional to the density of points at the value on the y axis. Horizontal black lines represent the 2.5%, 50%, and 97.5% quantiles.  $\overline{\Delta TSS}$ is the difference in TSS between the focal model and the corresponding model using the original 19 bioclim variables, calculated on a 10% random hold-out sample.

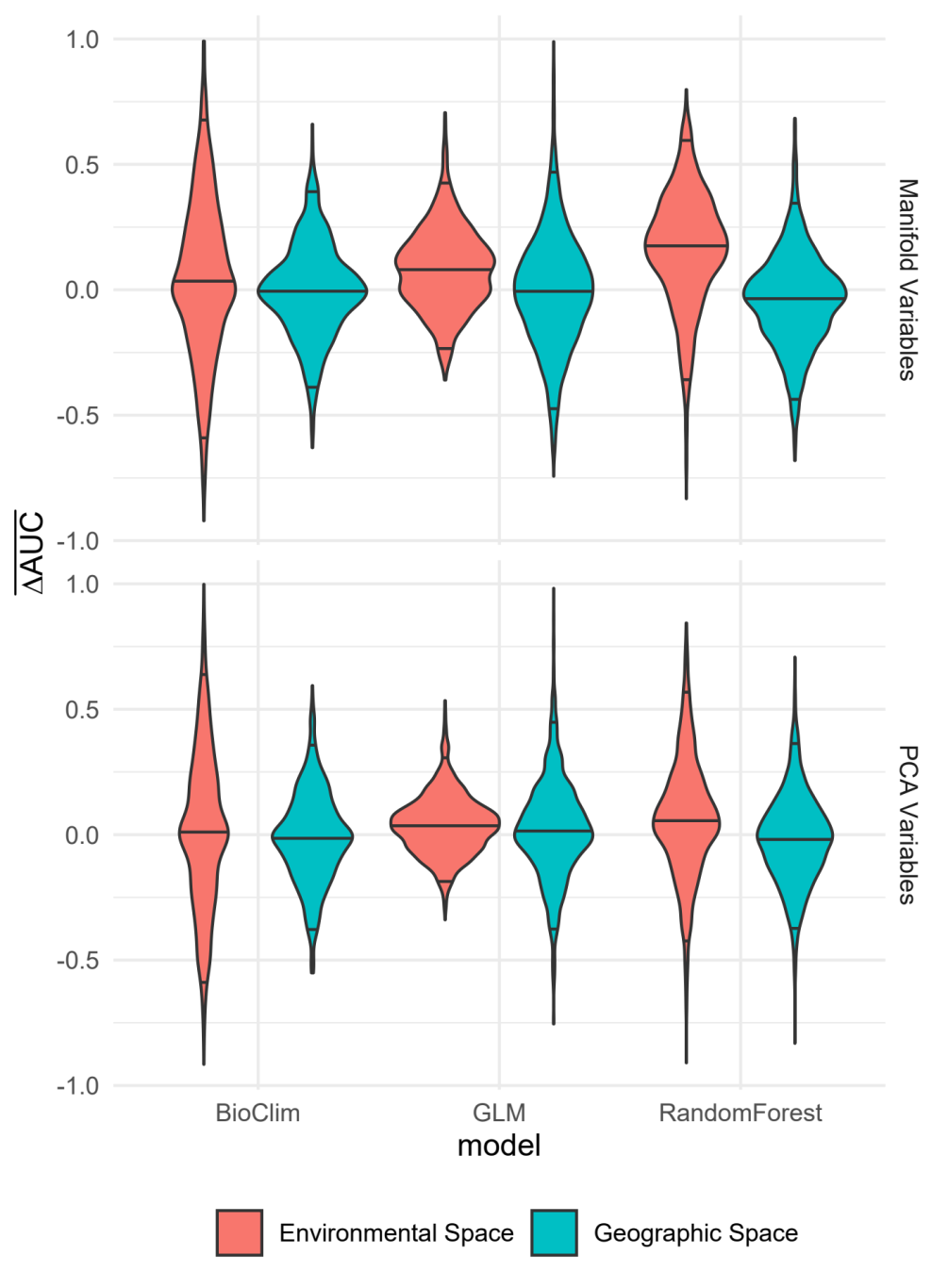

**Figure S3.** Violin plot of spatial block hold-out $\overline{\Delta AUC}$, by model type (X axis), Environmental vs. Geographic space (colour) and whether the models used the manifold or the PCA variables (top and bottom panels). Width of the violins is proportional to the density of points at the value on the y axis. Horizontal black lines represent the 2.5%, 50%, and 97.5% quantiles.  $\overline{\Delta AUC}$ is the difference in AUC between the focal model and the corresponding model using the original 19 bioclim variables, calculated on a spatial block hold-out sample.

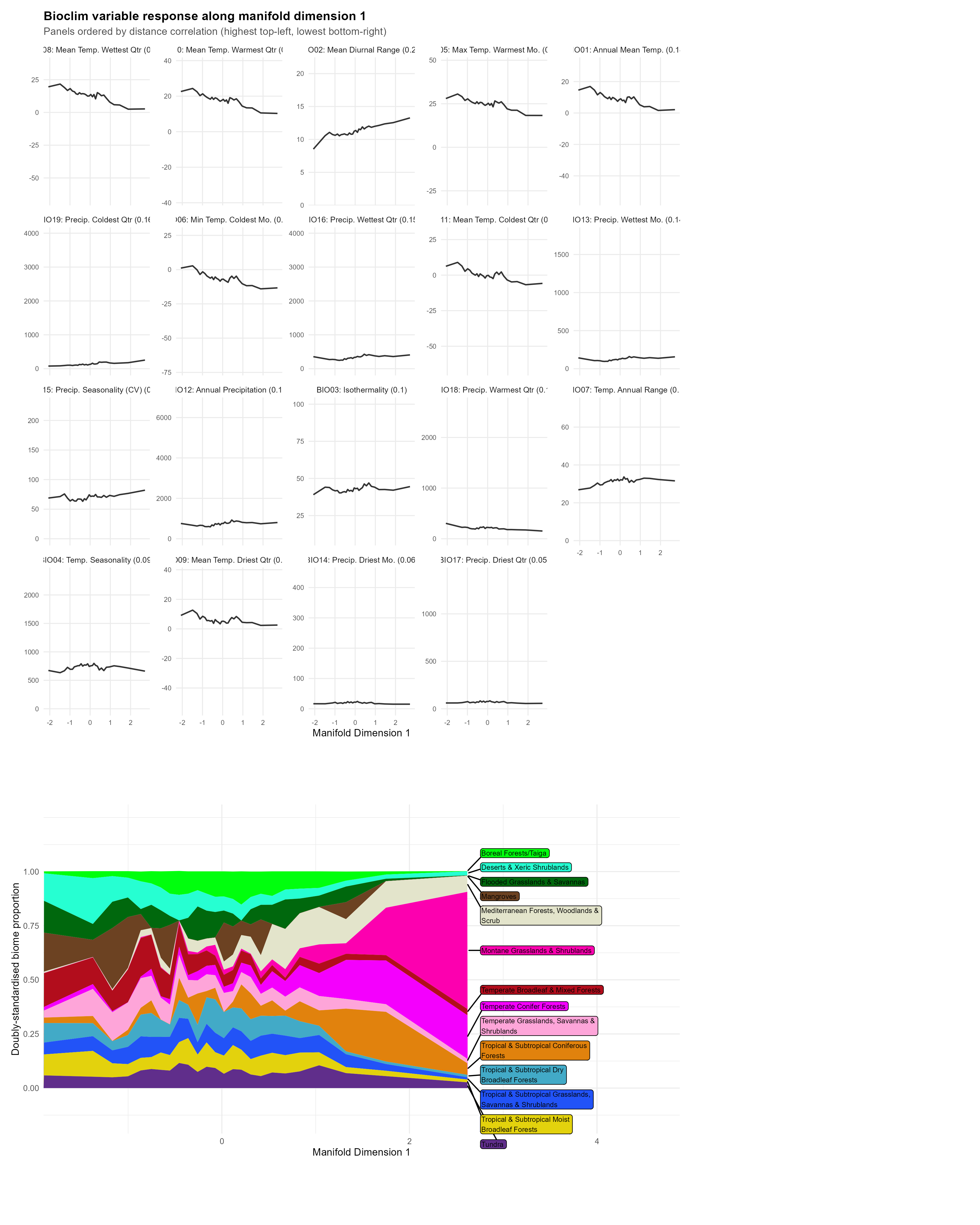

**Figure S4.** Examining and interpreting manifold dimension 1 of bioclim data. The top panel shows how the latent variable relates to the original 19 bioclim variables. Each bioclim variable is shown in a separate sub-panel, ordered by distance correlation with the manifold dimension (highest association top-left, lowest bottom-right). The line in each sub-panel shows the mean value of the bioclim variable (in original units) across equal-frequency bins of the manifold variable. The y-axis of each sub-panel spans the full range of the variable, so that weak relationships appear as flat lines. This gives an idea of how the manifold variable maps onto the original variables (similar to a non-linear version of loadings in PCA). The bottom panel shows how ecoregions are represented along the manifold variable. For each manifold variable bin the proportion of points that fell in each ecoregions was calculated and then doubly standardized by the total representation of each ecoregion globally. These data were then used to make a stacked area plot, where each colour-filled area corresponds to the proportion of the ecoregion along the manifold variable gradient.

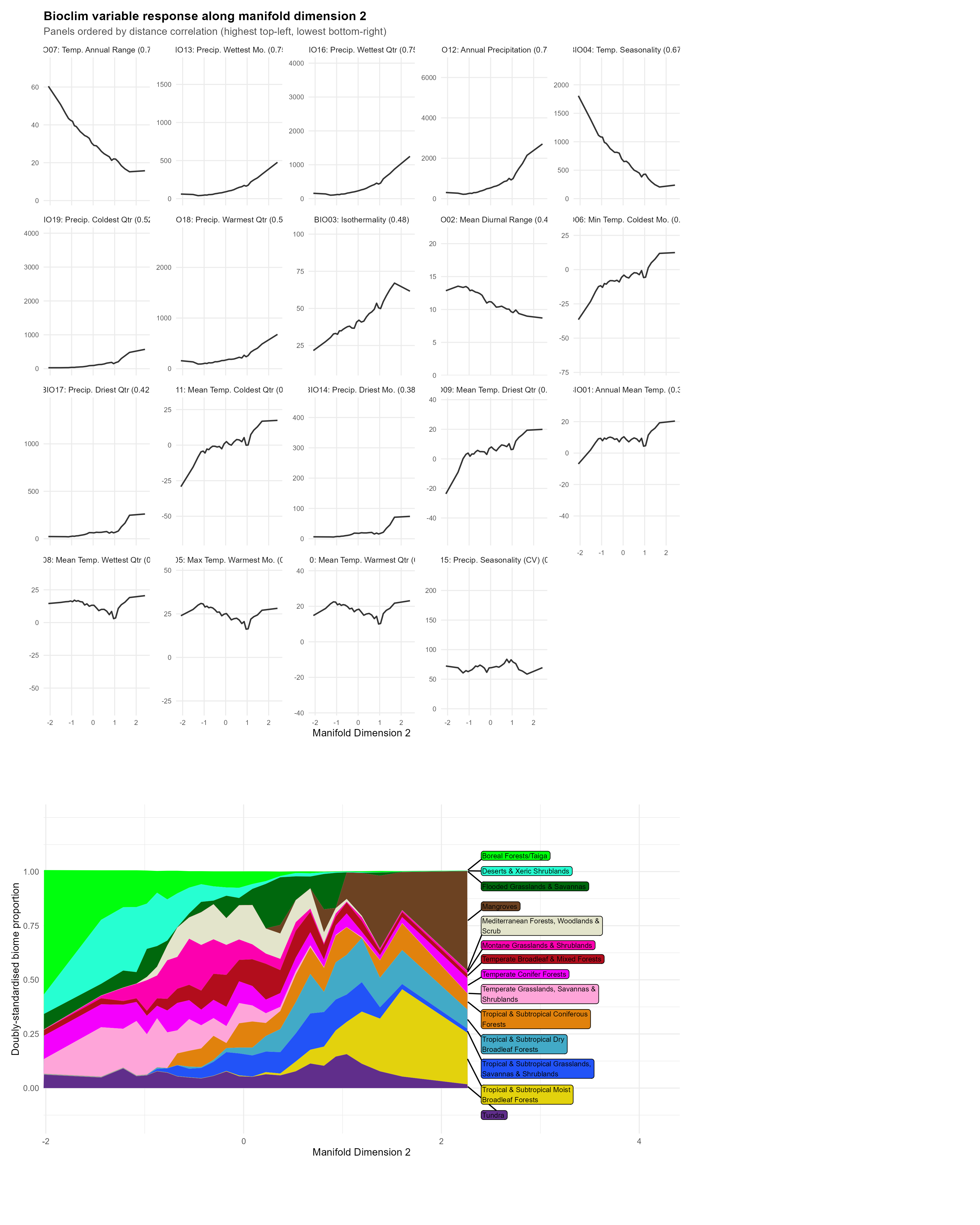

**Figure S5.** Examining and interpreting manifold dimension 2 of bioclim data.  The top panel shows how the latent variable relates to the original 19 bioclim variables. Each bioclim variable is shown in a separate sub-panel, ordered by distance correlation with the manifold dimension (highest association top-left, lowest bottom-right). The line in each sub-panel shows the mean value of the bioclim variable (in original units) across equal-frequency bins of the manifold variable. The y-axis of each sub-panel spans the full range of the variable, so that weak relationships appear as flat lines. This gives an idea of how the manifold variable maps onto the original variables (similar to a non-linear version of loadings in PCA). The bottom panel shows how ecoregions are represented along the manifold variable. For each manifold variable bin the proportion of points that fell in each ecoregions was calculated and then doubly standardized by the total representation of each ecoregion globally. These data were then used to make a stacked area plot, where each colour-filled area corresponds to the proportion of the ecoregion along the manifold variable gradient.

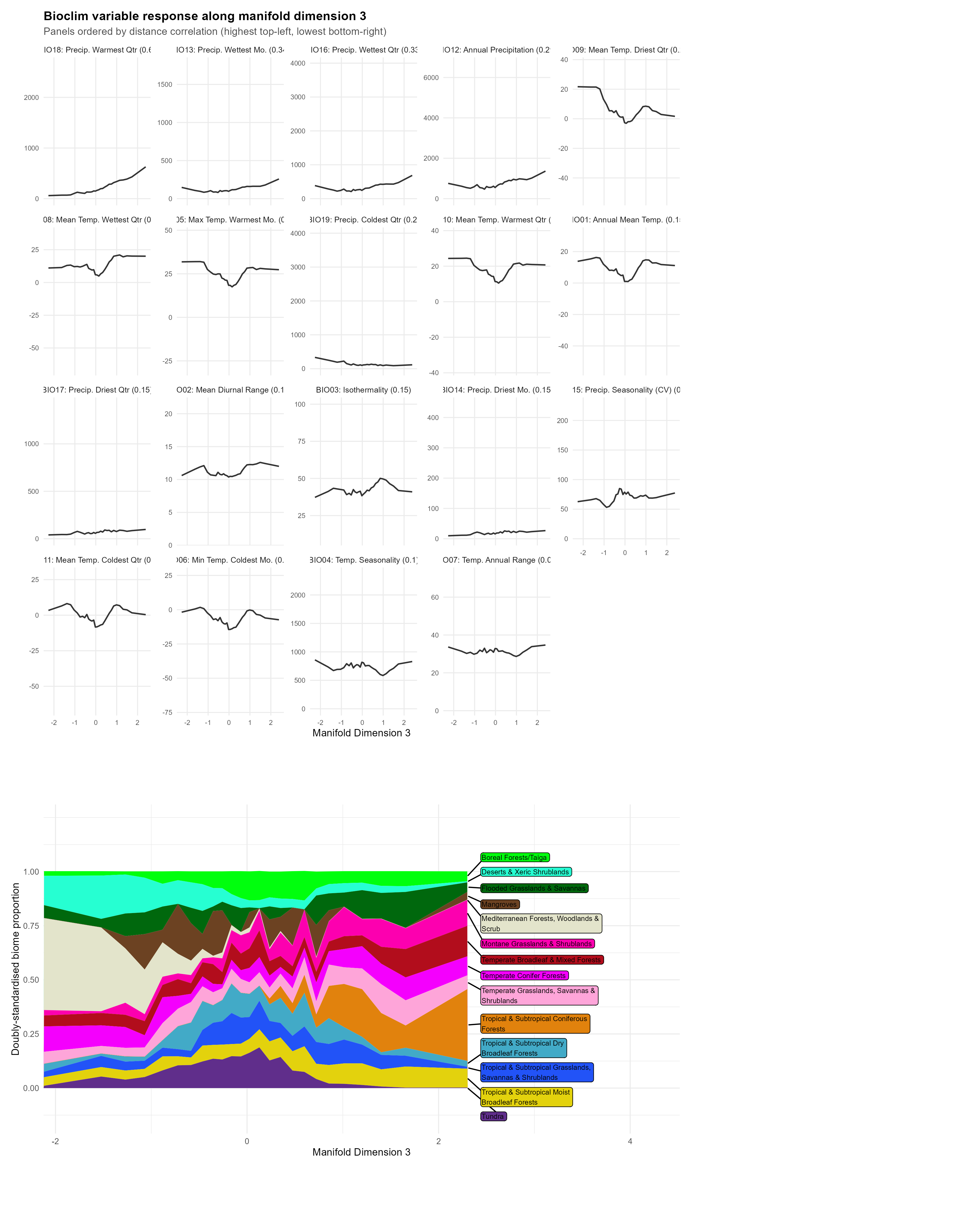

**Figure S6.** Examining and interpreting manifold dimension 3 of bioclim data.  The top panel shows how the latent variable relates to the original 19 bioclim variables. Each bioclim variable is shown in a separate sub-panel, ordered by distance correlation with the manifold dimension (highest association top-left, lowest bottom-right). The line in each sub-panel shows the mean value of the bioclim variable (in original units) across equal-frequency bins of the manifold variable. The y-axis of each sub-panel spans the full range of the variable, so that weak relationships appear as flat lines. This gives an idea of how the manifold variable maps onto the original variables (similar to a non-linear version of loadings in PCA). The bottom panel shows how ecoregions are represented along the manifold variable. For each manifold variable bin the proportion of points that fell in each ecoregions was calculated and then doubly standardized by the total representation of each ecoregion globally. These data were then used to make a stacked area plot, where each colour-filled area corresponds to the proportion of the ecoregion along the manifold variable gradient.

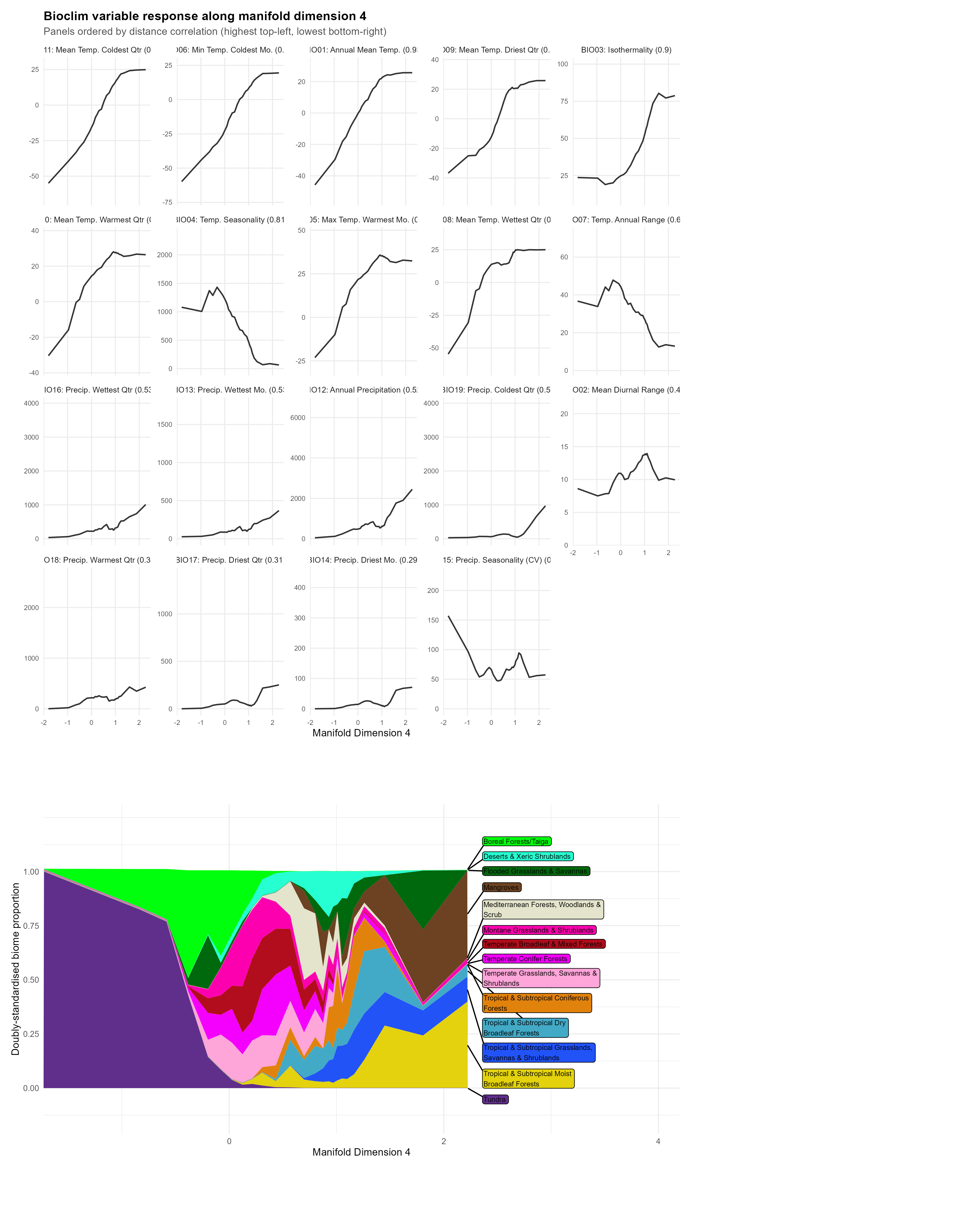

**Figure S7.** Examining and interpreting manifold dimension 4 of bioclim data. The top panel shows how the latent variable relates to the original 19 bioclim variables. Each bioclim variable is shown in a separate sub-panel, ordered by distance correlation with the manifold dimension (highest association top-left, lowest bottom-right). The line in each sub-panel shows the mean value of the bioclim variable (in original units) across equal-frequency bins of the manifold variable. The y-axis of each sub-panel spans the full range of the variable, so that weak relationships appear as flat lines. This gives an idea of how the manifold variable maps onto the original variables (similar to a non-linear version of loadings in PCA). The bottom panel shows how ecoregions are represented along the manifold variable. For each manifold variable bin the proportion of points that fell in each ecoregions was calculated and then doubly standardized by the total representation of each ecoregion globally. These data were then used to make a stacked area plot, where each colour-filled area corresponds to the proportion of the ecoregion along the manifold variable gradient.

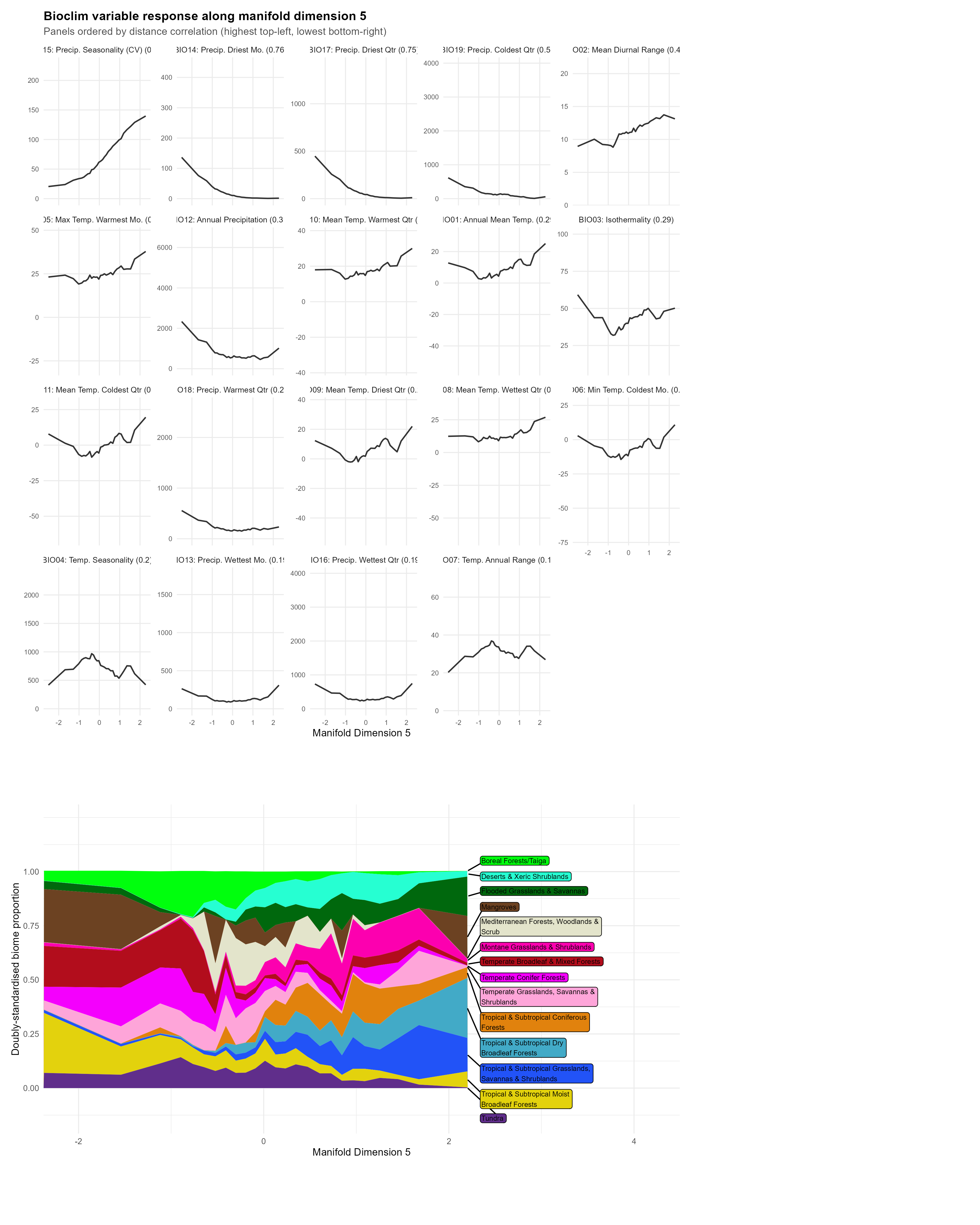

**Figure S8.** Examining and interpreting manifold dimension 5 of bioclim data. The top panel shows how the latent variable relates to the original 19 bioclim variables. Each bioclim variable is shown in a separate sub-panel, ordered by distance correlation with the manifold dimension (highest association top-left, lowest bottom-right). The line in each sub-panel shows the mean value of the bioclim variable (in original units) across equal-frequency bins of the manifold variable. The y-axis of each sub-panel spans the full range of the variable, so that weak relationships appear as flat lines. This gives an idea of how the manifold variable maps onto the original variables (similar to a non-linear version of loadings in PCA). The bottom panel shows how ecoregions are represented along the manifold variable. For each manifold variable bin the proportion of points that fell in each ecoregions was calculated and then doubly standardized by the total representation of each ecoregion globally. These data were then used to make a stacked area plot, where each colour-filled area corresponds to the proportion of the ecoregion along the manifold variable gradient.

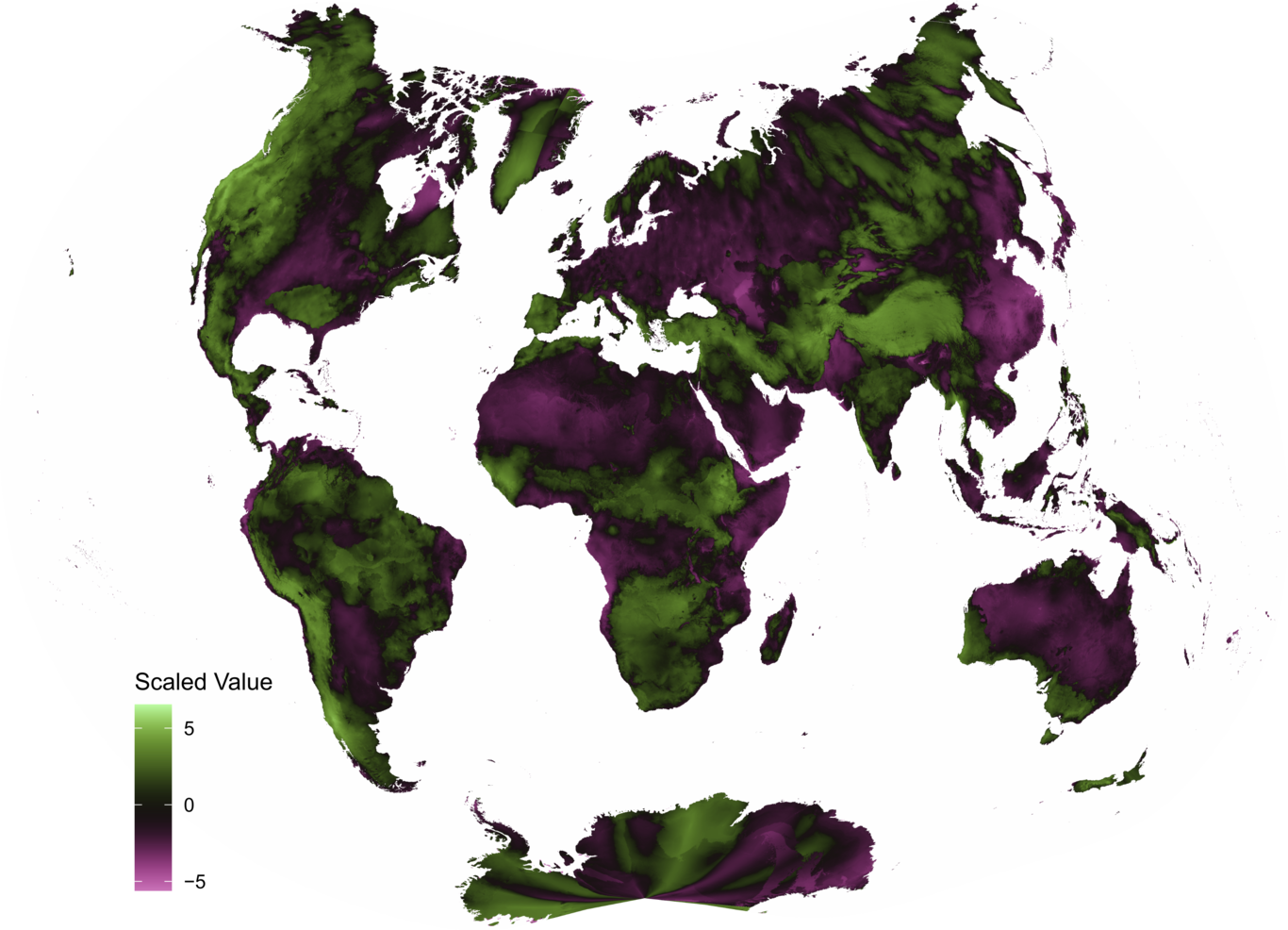

**Figure S9.** Map of bioclim manifold variable 1 in Daiseiji V projection (Kunimune 2020) .

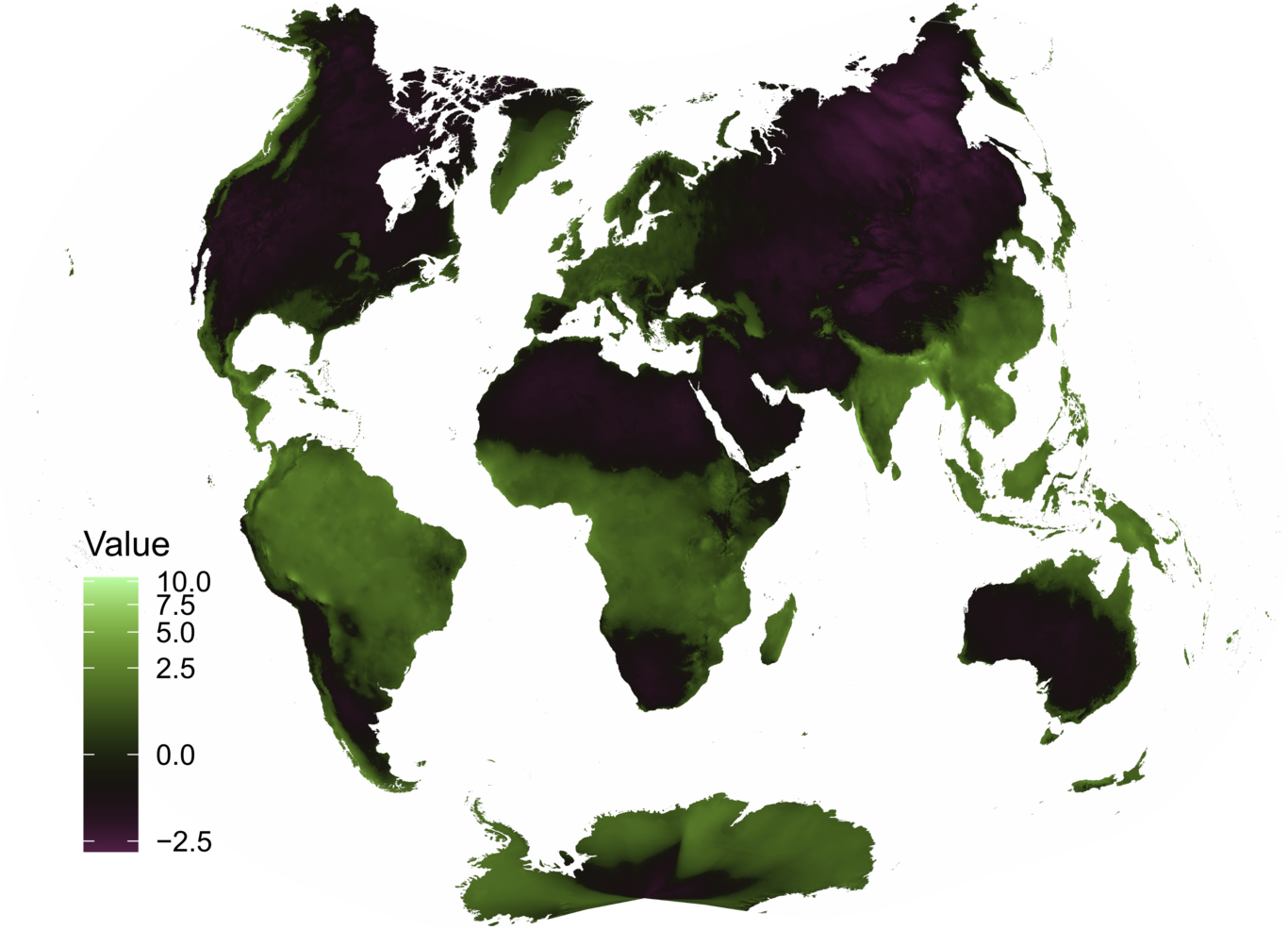

**Figure S10.** Map of bioclim manifold variable 2 in Daiseiji V projection  (Kunimune 2020).

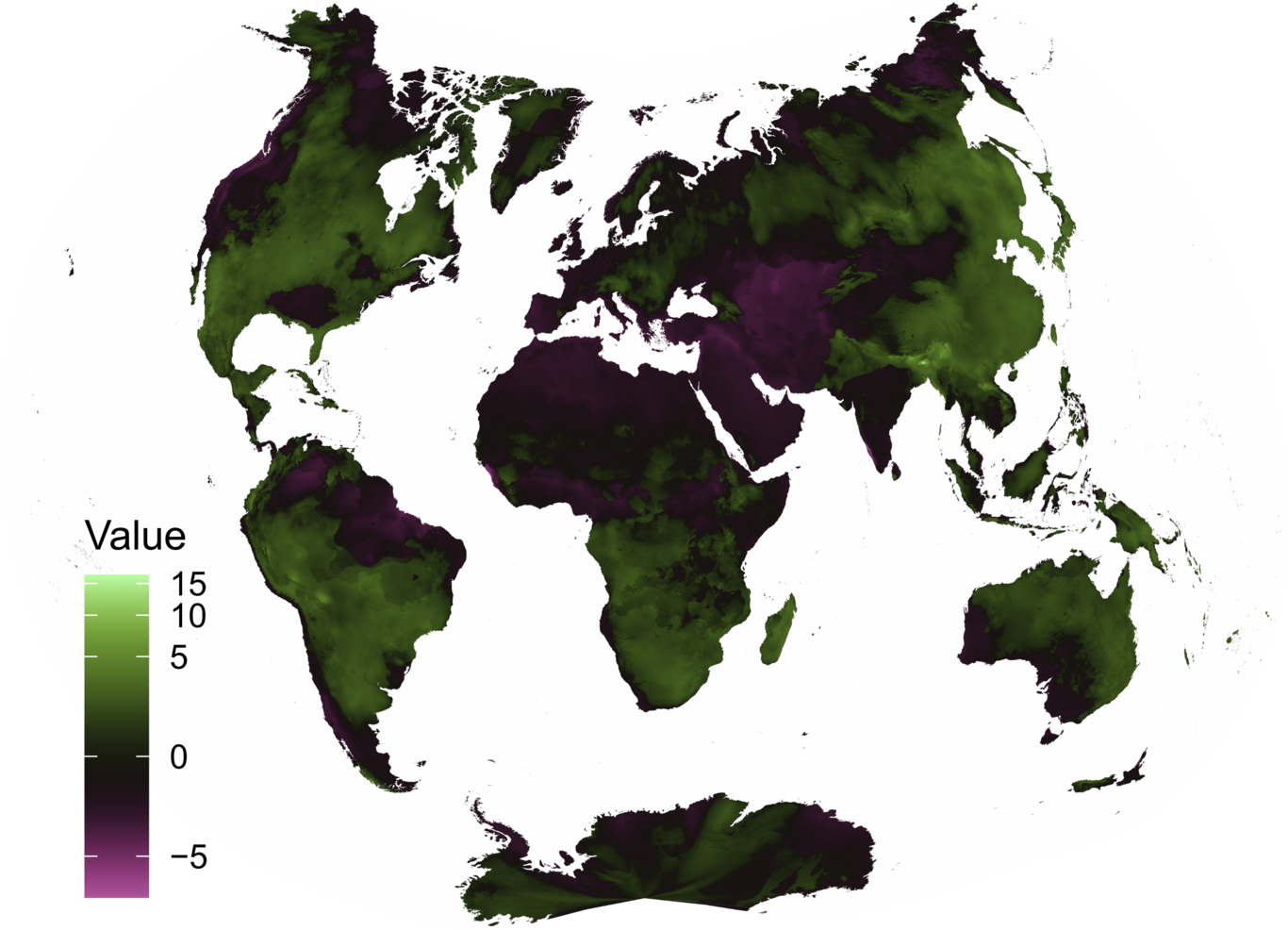

**Figure S11.** Map of bioclim manifold variable 3 in Daiseiji V projection  (Kunimune 2020).

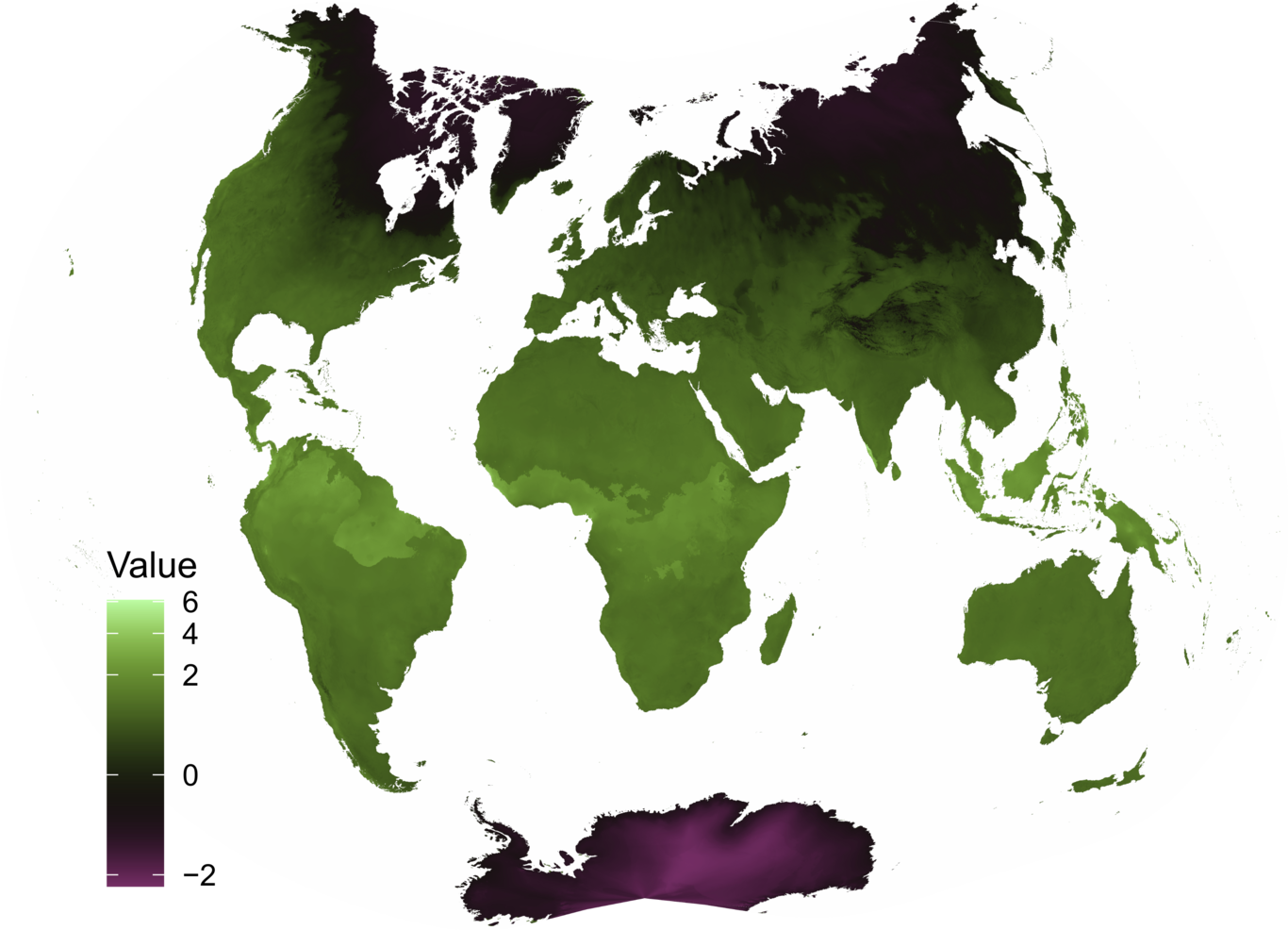

**Figure S12.** Map of bioclim manifold variable 4 in Daiseiji V projection  (Kunimune 2020).

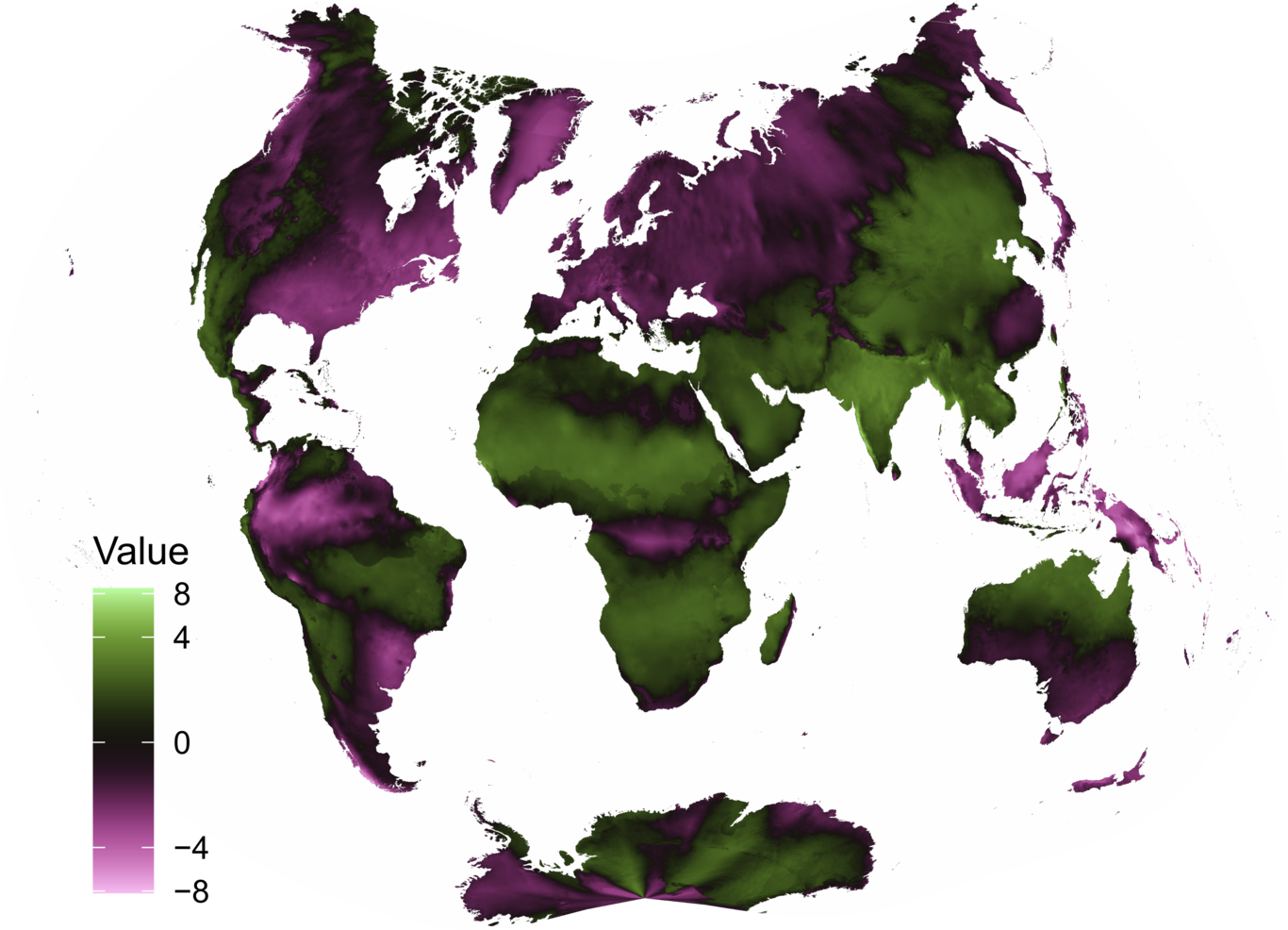

**Figure S13.** Map of bioclim manifold variable 5 in Daiseiji V projection  (Kunimune 2020).

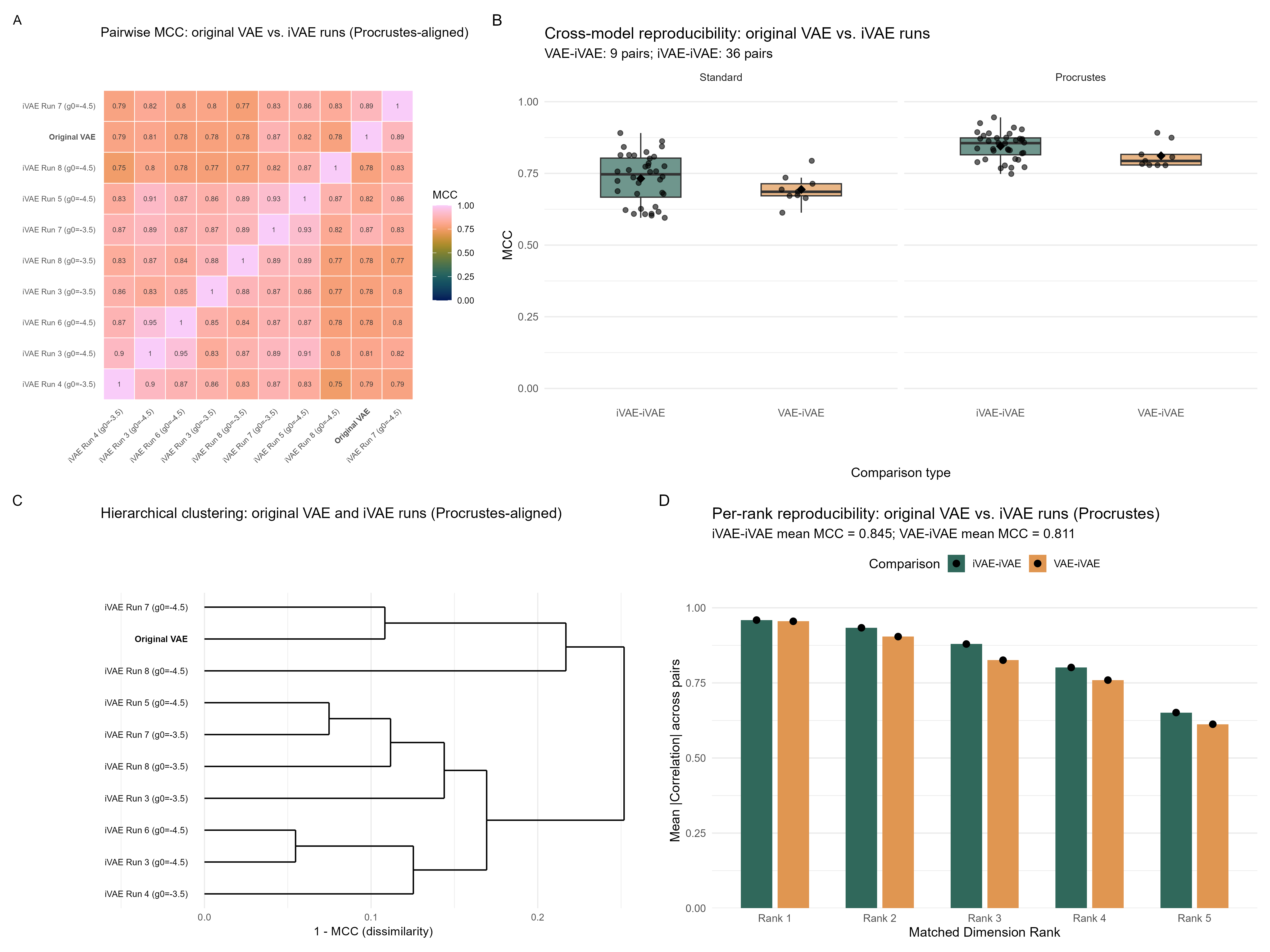

**Figure S14.** Comparison of the original (unconditional) VAE manifold with nine iVAE manifolds (Procrustes-aligned). (A) 10×10 MCC heatmap showing pairwise similarity between all runs (original VAE labelled in bold). (B) Distribution of MCC values by comparison type: iVAE-vs-iVAE (within-model) and VAE-vs-iVAE (cross-model), for both standard and Procrustes alignment. (C) Hierarchical clustering of all 10 runs by manifold dissimilarity (1 − MCC). (D) Per-rank reproducibility separated by comparison type, showing that the best-matched dimension (Rank 1) is recovered with mean |r| ≈ 0.96, while the weakest (Rank 5) has mean |r| ≈ 0.65.

#### Identifiability of VAE Manifold Dimensions

A known limitation of standard Variational Autoencoders (VAEs) is non-identifiability: the learned latent representations are not unique, meaning different training runs may recover different rotations of the same manifold subspace (Locatello et al. 2019). Identifiable VAEs (iVAEs; Khemakhem et al. 2020) address this by incorporating auxiliary observed information as a conditioning variable, which under certain conditions guarantees identifiability of the latent space up to a permutation and element-wise transformation. To assess whether the 5-dimensional manifold structure reported in the main text is robust to training variability, I conducted an identifiability analysis using iVAEs.

I trained 16 iVAE models using latitude and longitude, encoded as sinusoidal angle embeddings (described below), as the auxiliary conditioning variable. These comprised 8 random initialisations at each of two initial precision settings (log(γ₀) = −3.5 and −4.5). Of the 16 runs, 9 converged to exactly 5 active latent dimensions (defined as dimensions with mean posterior variance < 0.5), matching the original unconditional VAE. I restricted all subsequent comparisons to these 9 iVAE runs plus the original VAE (10 models total).

***iVAE Model Architecture and Training***

The identifiable VAE (iVAE) extends the standard VAE framework by introducing a conditional prior on the latent variables that depends on an observed auxiliary variable u. While a standard VAE assumes a fixed prior p(z) = N(0, I), the iVAE learns a conditional prior p(z|u) parameterised by a neural network, where u provides additional observed context for each data point. Khemakhem et al. (2020) showed that this conditioning, combined with sufficient variability in the auxiliary variable, guarantees identifiability of the latent representations up to permutation and element-wise transformation. The full iVAE model is implemented in R using the torch package (Falbel and Luraschi 2022) and the dagnn package (Dinnage, in development) for specifying neural network architectures as directed acyclic graphs with skip connections. The complete source code is available at https://github.com/rdinnager/bioclim_intrinsic_dimension.

Sinusoidal angle embedding. Geographic coordinates (latitude and longitude in decimal degrees) were encoded into a 64-dimensional vector using sinusoidal angle embeddings, with 32 dimensions allocated to each coordinate. For each coordinate value θ (in the original degree units), the embedding applies a set of 16 frequency bands fₖ = 2⁰, 2¹, 2², …, 2¹⁵, producing interleaved sine and cosine pairs: [sin(f₁θ), cos(f₁θ), sin(f₂θ), cos(f₂θ), …, sin(f₁₆θ), cos(f₁₆θ)]. This encoding is analogous to positional encodings in transformer models (Vaswani et al. 2017) and provides a smooth, continuous representation that captures geographic relationships at multiple spatial scales. The lowest frequency captures broad hemispheric patterns, while the highest frequencies resolve differences of fractions of a degree. The latitude and longitude embeddings were concatenated to form the 64-dimensional auxiliary variable u used by the iVAE encoder and prior network.

Network architecture. The iVAE comprises three neural network components: an encoder, a decoder, and a prior network, each implemented as a three-hidden-layer feedforward network with ReLU activations and skip connections via nndag(). The encoder takes as input the 19 standardised bioclimatic variables concatenated with the 64-dimensional angle embedding u, and outputs the mean and log-variance of the approximate posterior q(z|x, u) over 16 latent dimensions. Each hidden layer has 32 units (2 × latent dimensions) and receives a skip connection from the angle embedding. The prior network takes only the angle embedding u as input and outputs a location-dependent prior distribution p(z|u), parameterised by its own mean and log-variance vectors over the same 16 latent dimensions. The decoder takes only the latent sample z (without the conditioning variable) and outputs the reconstructed 19 bioclimatic variables. The decoder’s independence from the conditioning variable is important: it ensures that the latent representation z must capture all information needed to reconstruct the bioclimatic variables, while the prior network modulates the expected latent distribution based on geographic location.

Loss function. The model is trained by maximising a modified Evidence Lower Bound (ELBO). The loss function comprises three terms: (1) a reconstruction loss measuring the squared error between input and reconstructed bioclimatic variables, scaled by a learnable precision parameter γ (following Dai and Wipf 2019); (2) the KL divergence between the approximate posterior q(z|x, u) and the conditional prior p(z|u), which encourages the encoder to produce representations consistent with the location-dependent prior; and (3) a weighted KL divergence between the conditional prior p(z|u) and a standard normal distribution N(0, I), weighted by a hyperparameter (set to 0.1), which regularises the prior network and prevents it from collapsing to a degenerate distribution. The learnable precision parameter γ (parameterised as log(γ)) automatically balances reconstruction fidelity against latent space regularisation during training. As in the original VAE, latent dimensions whose mean posterior variance exceeds 0.5 across the dataset are considered inactive and pruned, allowing the model to automatically determine the intrinsic dimensionality.

Training details. The 19 bioclimatic variables at 2.5 arc-minute resolution were standardised to zero mean and unit standard deviation before input. Each iVAE model was trained for 5,000 epochs using the Adam optimiser (Kingma and Ba 2015) with a learning rate of 0.0005 and a one-cycle learning rate schedule (Smith and Topin 2019). Batch size was 1,300,000 data points per batch. The log-precision parameter log(γ) was initialised to either −3.5 or −4.5 across the two sets of runs. Training was performed on an NVIDIA GPU using the torch package for R. All 16 runs used the same data but different random initialisations of the network weights, providing a test of the stability and identifiability of the learned representations.

To quantify the similarity of learned representations across runs, I computed the pairwise Mean Correlation Coefficient (MCC). The MCC uses the Hungarian algorithm to find the optimal one-to-one matching of latent dimensions that maximises the mean absolute Pearson correlation between matched dimension pairs. Because different training runs may learn the same subspace but express it through different rotations, I also computed the MCC after Procrustes alignment, in which an SVD-based orthogonal rotation is applied to align one run’s latent space to another’s before matching dimensions.

Without Procrustes alignment, the mean pairwise MCC across all 45 pairs (10 models choose 2) was 0.724, indicating substantial but imperfect dimension-by-dimension correspondence. After Procrustes alignment, the mean MCC increased to 0.838, confirming that different runs recover the same subspace but may express it through different rotations. The iVAE-vs-iVAE Procrustes MCC was 0.845 (sd = 0.048, range: 0.748–0.945), while the VAE-vs-iVAE Procrustes MCC was 0.811 (sd = 0.043, range: 0.778–0.892). A hierarchical clustering dendrogram of pairwise dissimilarities (1 − MCC) showed that the original VAE clusters among the iVAE runs rather than as an outgroup, indicating that the unconditional VAE recovers essentially the same manifold structure as the identifiable models.

These results demonstrate that the 5-dimensional manifold structure is a robust and reproducible feature of the bioclimatic data, rather than an artifact of a particular random initialisation. The high agreement between the original VAE and the iVAE runs indicates that the manifold dimensions used in the SDM comparisons are not an idiosyncratic product of a single training run.

Not all manifold dimensions were equally reproducible across runs. When matched dimension pairs are ranked from best to worst correlation for each pairwise comparison, the best-matched dimension (Rank 1) was recovered with a mean absolute correlation of 0.96 across all comparisons, indicating near-perfect reproducibility. Subsequent ranks showed progressively lower correlations: Rank 2 (mean |r| ≈ 0.93), Rank 3 (mean |r| ≈ 0.88), Rank 4 (mean |r| ≈ 0.79), and Rank 5 (mean |r| ≈ 0.65). This gradient was similar for both iVAE-vs-iVAE and VAE-vs-iVAE comparisons (Figure S14D), suggesting that the ordering of dimensions by identifiability is a property of the data rather than the model type. The most reproducible dimensions likely correspond to the strongest climatic gradients (e.g. the precipitation and temperature axes captured by Manifold dimensions 1 and 3), while the least reproducible dimension may capture subtler variation that is more sensitive to training conditions.
